# Gut microbiome composition better reflects host phylogeny than diet in breeding wood-warblers

**DOI:** 10.1101/2022.03.07.482310

**Authors:** Marcella D. Baiz, Andrea Benavides C., Eliot T. Miller, Andrew W. Wood, David P. L. Toews

## Abstract

Understanding the factors that shape microbiomes can provide insight on the importance of host-symbiont interactions and on co-evolutionary dynamics. Unlike for mammals, previous studies have found little or no support for an influence of host evolutionary history on avian gut microbiome diversity and instead have suggested a greater influence of the environment or diet due to fast gut turnover. Because effects of different factors may be conflated by captivity and sampling design, examining natural variation using large sample sizes is important. Our goal was to overcome these limitations by sampling wild birds to compare environmental, dietary, and evolutionary influences on gut microbiome structure. We performed fecal metabarcoding to characterize both the gut microbiome and diet of fifteen wood-warbler species across a four-year period and from two geographic localities. We find host taxonomy generally explained ∼10% of the variation between individuals, which is ∼6-fold more variation of any other factor considered, including diet diversity. Further, gut microbiome similarity was more congruent with the host phylogeny than with host diet similarity and we found little association between diet diversity and microbiome diversity. Together, our results suggest evolutionary history is the strongest predictor of gut microbiome differentiation among wood-warblers. Although the phylogenetic signal of the warbler gut microbiome is not very strong, our data suggest that a stronger influence of diet (as measured by diet diversity) does not account for this pattern. The mechanism underlying this phylogenetic signal is not clear, but we argue host traits may filter colonization and maintenance of microbes.

## Introduction

Microorganisms that form intimate associations with their hosts can take part in important physiological functions. In particular, the gut microbiome—the community of microbes that colonize the gastrointestinal tract—has been linked to host behavior, immune function, metabolism, and disease (Sommer & Backhed 2013, Suzuki 2017, Bodawatta et al. 2021b).

The taxonomic composition of the gut microbiome can vary, sometimes dramatically within and between host species (Loo et al. 2019, Grond et al. 2019, Song et al. 2020), as well as within-individuals over short timescales (Videvall et al. 2019, Skeen et al. 2021). However, when host-microbe associations are long-term, gut microbiomes may be expected to be species-specific and their assembly to be dependent on host evolutionary divergence (Brooks et al. 2016). Consistent with this, host evolutionary history, in addition to diet, has been implicated as one of the strongest factors driving vertebrate gut microbiome similarity (Youngblut et al. 2020).

Recent studies have strongly supported a positive correlation between host species divergence and gut microbiome divergence—known as “phylosymbiosis”—particularly for insects and non-flying mammals (Brooks et al. 2016, Song et al. 2020). However, in birds, differences in gut microbiome structure between species are less pronounced (Song et al. 2020). Despite species-level differences in gut microbiota of 37 New Guinean passerine species (14 families), Bodawatta et al. (2021a) did not find an influence of host phylogeny on gut microbiome structure. This is in contrast, however, to a study on 51 passerine species (21 families) breeding in the Czech Republic (Kropáková et al. 2017) and a study on all 15 crane species (family Gruidae) in captivity which found a weak influence of host phylogeny, and only when examining female individuals (Trevelline et al. 2020).

A favorable hypothesis to explain this marked difference in phylosymbiosis between bird and non-flying mammal gut microbiota is that because birds evolved a reduced and simplified gastrointestinal tract as an adaptation to flight, they have highly reduced gut retention times from consumption of food to defecation (Song et al. 2020). This reduced retention time and simplified gut environment may favor high turn-over in the avian gut microbiome, and a larger role of the diet and environment over host taxonomy in the structuring of the gut microbiome (Bodawatta et al. 2021a).

In Darwin’s finches, gut microbiome communities cluster more strongly by host habitat than by host species (Loo et al. 2019). Host phylogeny and diet in this group, which is known for adaptive divergence in beak morphology that is linked to foraging ecology, both show a moderate influence on gut microbiome variation (Loo et al. 2019). Further, the gut microbiome of the vampire finch, a diet specialist, is highly divergent from other species (Michel et al. 2018). Similarly, captive birds tend to have distinct gut microbiota from their wild counterparts (San Juan et al. 2021). These studies support a strong role of the environment, including diet, in shaping the avian gut microbiome.

Although many studies have detected effects of diet on the avian gut microbiome (Xiao et al. 2021, Bodawatta et al. 2021a, Davidson et al. 2020, Knutie 2020, Teyssier et al. 2020), few have analyzed host diet beyond broad categorizations of diet type (e.g., omnivore versus insectivore) and/or included birds that were fed standardized and non-natural diets (but see Bodawatta et al. 2022, Schmiedová et al. 2022). Further, many studies that have assessed species-specific differences in gut microbiome structure have had limited sample sizes including only one or a few individuals per species or included data collected and sequenced at different times or in different ways. To gain a holistic picture of the effects of host diet, evolutionary history, and geography on gut microbiome structure, it will be necessary to sample natural populations using standardized methods. Understanding the factors that shape the avian gut microbiome is important for understanding host-symbiont interactions and co-evolutionary dynamics, and how these dynamics may differ from other taxonomic groups of animals (i.e., mammals). The role of the gut microbiome in host evolutionary processes is largely unexplored and its potential role in facilitating and responding to avian host adaptive radiation—where species diversification is tied to ecological differentiation—is a major outstanding question (Bodawatta et al. 2021b).

Here, we characterize the gut microbiome of wood-warblers (family: Parulidae) breeding in sympatry in Eastern North America across a 4-year period and examine factors that may play a role in shaping gut microbiome structure. Parulidae is a passerine radiation of >100 insectivorous species that evolved rapidly in the last 7 MY (Lovette et al. 2010, Barker et al. 2015), and is a classic model for studies of ecological differentiation, including diet niche partitioning (MacArthur 1958). In the current study, we use 16S fecal metabarcoding to examine gut microbiomes of 15 species representing 7 genera (Figure 1a). Our aims are to characterize the “core” parulid gut microbiota (a common set of microbes across individuals) and to quantify differences in gut microbiome composition between hosts. We predict that due to genetic and ecological differentiation among host species, variation in the gut microbiome will be largely explained by host taxonomy. Further, we explicitly test the prediction of phylosymbiosis, where host phylogenetic relatedness should correlate with gut microbiome similarity. We also examine the relationship between gut microbiome diversity and diet diversity by analyzing COI metabarcoding sequences amplified from fecal samples of these same individuals. With the presumption that a diet characterized by a high diversity of arthropods will incur ingestion of a greater diversity of bacteria—either associated with arthropod hosts, or the environments in which they are found—we predict that diversity of the warbler gut microbiome and diet will be positively correlated. Finally, we test for other environmental signals in the structuring of host gut microbiomes by examining effects of sampling year, locality, and diet specialization.

**Figure 1.**
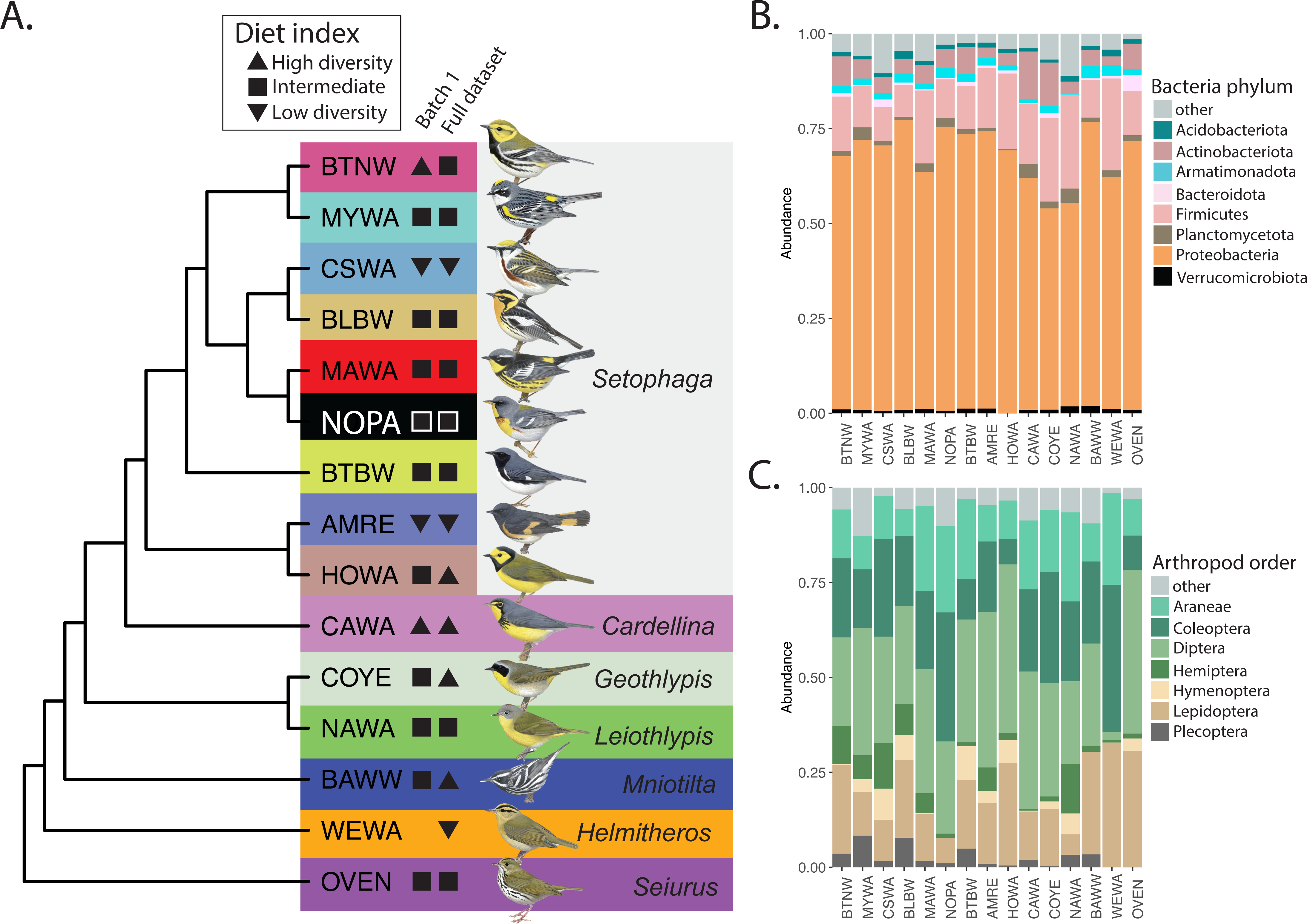
**A)** Phylogenetic relationships between host species in this study. Upside down triangles indicate low diversity diet, triangles indicate high diversity diet, and squares indicate intermediate diet diversity based on our COI diet index. The full dataset represents all samples collected between 2017-2020, and batch 1 represents all samples collected between 2017-2019. Illustrations © Lynx Edicions. **B)** Relative abundance of bacterial phyla in the full 16S dataset, and **C)** Relative abundance of arthropod orders in the full COI dataset. See Table 1 for host species codes.

**Table 1.** Sampling information for species included in this study. NY=New York, PA=Pennsylvania. Subset batch 1 includes samples collected between 2017-2019, which were sequenced together in a single sequencing run.

| Species code | English common name | Latin name | 2017 |  | 2018 |  | 2019 |  | 2020 |  | Total |  |
| --- | --- | --- | --- | --- | --- | --- | --- | --- | --- | --- | --- | --- |
|  |  |  | NY | PA | NY | PA | NY | PA | NY | PA | Subset batch 1 | Full dataset |
| AMRE | American Redstart | *Setophaga ruticilla* | 2 | 0 | 3 | 0 | 5 | 6 | 4 | 4 | 16 | 23 |
| BAWW | Black-and-white Warbler | *Mniotilta varia* | 0 | 0 | 7 | 0 | 1 | 1 | 4 | 9 | 9 | 22 |
| BLBW | Blackburnian Warbler | *Setophaga fusca* | 1 | 0 | 4 | 0 | 3 | 1 | 5 | 6 | 9 | 20 |
| BTBW | Black-throated Blue Warbler | *Setophaga caerulescens* | 2 | 0 | 4 | 0 | 2 | 5 | 5 | 10 | 13 | 28 |
| BTNW | Black-throated Green Warbler | *Setophaga virens* | 3 | 0 | 4 | 0 | 4 | 3 | 5 | 7 | 14 | 26 |
| CAWA | Canada Warbler | *Cardellina canadensis* | 2 | 0 | 4 | 0 | 3 | 3 | 4 | 1 | 12 | 17 |
| COYE | Common Yellowthroat | *Geothlypis trichas* | 1 | 0 | 4 | 0 | 3 | 1 | 3 | 2 | 9 | 14 |
| CSWA | Chestnut-sided Warbler | *Setophaga pensylvanica* | 5 | 0 | 2 | 0 | 1 | 5 | 3 | 3 | 13 | 18 |
| HOWA | Hooded Warbler | *Setophaga citrina* | 0 | 0 | 0 | 0 | 0 | 6 | 0 | 5 | 6 | 11 |
| MAWA | Magnolia Warbler | *Setophaga magnolia* | 1 | 0 | 6 | 0 | 4 | 1 | 4 | 1 | 12 | 17 |
| MYWA | Myrtle Warbler | *Setophaga coronata* | 4 | 0 | 5 | 0 | 5 | 1 | 5 | 0 | 15 | 20 |
| NAWA | Nashville Warbler | *Leiothlypis ruficapilla* | 2 | 0 | 4 | 0 | 5 | 0 | 0 | 0 | 11 | 11 |
| NOPA | Northern Parula | *Setophaga americana* | 2 | 0 | 4 | 0 | 5 | 0 | 5 | 0 | 11 | 16 |
| OVEN | Ovenbird | *Seiurus aurocapilla* | 2 | 0 | 4 | 0 | 4 | 1 | 5 | 6 | 11 | 22 |
| WEWA | Worm-eating Warbler | *Helmitheros vermivorum* | 0 | 0 | 0 | 0 | 0 | 0 | 0 | 5 | 0 | 5 |
| Total individuals |  |  |  |  |  |  |  |  |  |  | 161 | 270 |

## Materials and Methods

### Sample collection and DNA extraction

We used mist nets to capture birds during four consecutive breeding seasons (May-July 2017-2020). In all years, we targeted sampling locations in northern hardwood forests, both in Adirondack Park, New York, and in 2019 and 2020, we also sampled birds in central Pennsylvania (Figure S1, Table 1). We selected sites where a diversity of warbler species (up to eight) could be heard singing so as to maximize sympatry among species included in the study. Upon capture, we held individuals inside a brown paper bag for up to 10 minutes to allow ample time for excretion inside the bag before removal and subsequent banding. We removed feces by scraping it from the inside of the bag directly into a sample tube containing lysis buffer (100 mM Tris pH 8; 100 mM Na _2_ EDTA, 10 mM NaCl; 0.5% sodium dodecyl sulfate; White & Densmore 1992), and froze samples at -20 °C within two weeks of collection. Because we were interested in variation among individuals, we chose a single sample at random to include in our analyses from individuals that were recaptured in the same or subsequent years. In total, we sequenced samples from 408 individuals.

We extracted total DNA from fecal samples using an SPRI-bead fecal DNA extraction method modified from Vo & Jedlicka (2014). Samples were processed in two sets: those collected in 2017-2019 and in 2020. After thawing fecal samples at room temperature, we centrifuged sample tubes and used bleach-sterilized laboratory spatulas after being thoroughly dried and/or pipetting to transfer ∼5 mg of fecal material into 2mL screw-cap microcentrifuge tubes each containing 0.25g of 0.1mm and 0.25g of 0.5mm zirconia-silica beads. For samples that amounted to <5 mg of fecal material, we supplemented with a suitable volume of storage buffer from inside the sample tube as necessary. We immediately added 818 μL of warmed (65°C) lysis buffer (Vo & Jedlicka 2014) and homogenized samples using a Precellys 24 Tissue Homogenizer (Bertin Instruments) set to 3 cycles of 6800rpm for 30s with a 30s pause between cycles. After transferring the supernatant to clean microfuge tubes, we incubated samples with Qiagen Solution C3 (Qiagen DNeasy PowerSoil 12888-100-3) to remove PCR inhibitors. Next, we removed DNA from the supernatant using homemade solid phase reversible immobilization (SPRI) magnetic beads (“Serapure” beads) (Rohland & Reich 2012). Serapure beads were added at a 1.9x bead-to-supernatant volume ratio and, after cleaning with 80% ethanol, we eluted DNA in 10mM Tris-HCL. Extracted DNA was stored at -20°C before proceeding with library preparation. We also included negative extraction controls that followed the same procedure described above for which the input was sample storage buffer taken from tubes that were transported to the field, but were not used for collecting fecal material.

### *16S* and COI amplicon sequencing

As with DNA extractions, we prepared and sequenced metabarcoding libraries in two separate batches: (1) samples collected between 2017-2019, and (2) samples collected in 2020. We used a two-step multiplex dual-index amplicon approach to separately prepare 16S libraries and COI libraries for sequencing again following Vo & Jedlicka (2014) with some adjustments. We first used universal 515F/806R primers to amplify the V4 region of the bacterial 16S rRNA gene (Caporaso et al. 2012) and the “ANML” general arthropod COI mitochondrial primers LCOI-1490/COI-CFMRa described in Jusino et al. (2019). Each primer pair was modified with overhanging Illumina adapter sequences. Prior to PCR, we randomized the order of samples to be amplified to avoid within-plate batch effects during amplification. Negative PCR controls were included on each plate. In addition to our fecal samples, we sequenced four negative controls per primer pair in each library pool, with the exception of the first batch COI library pool which did not contain any negative controls. Negatives included two “extraction controls” amplified and sequenced from DNA extractions made from sample tubes containing only buffer (and no feces) as well as two negative PCR controls.

We performed initial 16S PCR amplification for each sample in triplicate in 30 μL reactions comprising 0.2 μL Platinum II *Taq* Hot Start DNA Polymerase (Invitrogen 14966005), 5 μL 5X Buffer (Invitrogen 14966005), 1.25 μL of each primer (10uM concentration), 13.5 μL molecular grade water, and 0.5 μL 10mM dNTP mix (Promega U151A) and 3.3 μL of fecal DNA. Reaction conditions followed the 2-step PCR protocol recommended by the manufacturer: 94°C for 2m, followed by 34 cycles of 98°C for 5s, 68°C for 15s, followed by a final extension at 68°C for 5m, and hold at 12°C. We performed initial COI PCR amplification in 30 μL reactions comprising 0.24 μL Platinum II *Taq* Hot Start DNA Polymerase, 6 μL 5X Buffer, 1.5 μL of each primer (10uM concentration), 16.16μL molecular grade water, and 0.6 μL 10mM dNTP mix, and 4 μL of fecal DNA. Reaction conditions followed Jusino et al. (2019) with minor adjustments: 94°C for 2m, followed by 5 cycles of 94°C for 15s, 45°C for 15s, 68°C for 15s, followed by 35 cycles of 98°C for 5s, 68°C for 15s, followed by a final extension at 68°C for 5m, and hold at 12°C. We cleaned initial PCR products by incubating with a 1x volume of serapure beads and eluting the bound DNA in 10mM Tris-HCL. Triplicate 16S reactions were pooled before this cleaning step. Then we evaluated amplification success by visualizing cleaned product on a 1.5% agarose gel.

Next, we appended dual P5 and P7 Illumina indexes to each library via PCR. Reactions were 30 μL and contained 15 μL KAPA HiFi HotStart ReadyMix (Roche 7958935001), 3 μL of each primer (10uM concentration), and 9 μL DNA (cleaned initial PCR product). Reaction conditions followed manufacturer recommendations: 98°C for 45s, followed by 7 cycles of 98°C for 15s, 60°C for 15s, 72°C for 15s, followed by a final extension at 72°C for 1m, and hold at 12°C. We then cleaned the indexed PCR product using a double-sided serapure bead procedure. We first removed potential high-molecular weight contamination by incubating PCR product with a 0.75x volume of serapure beads. After placing the samples on the magnet, we transferred the supernatant to fresh tubes and incubated it with a 1x volume of serapure beads to remove potential low-molecular weight contamination. DNA was eluted in 10mM Tris-HCL, and we evaluated amplification success as for the initial PCR.

We quantified DNA in our final PCR products with a Qubit 4.0 Fluorometer (Invitrogen). We then normalized library concentrations and pooled libraries to a final pool concentration of at least 2nM. We submitted the final pool to the Penn State Genomics Core Facility to perform final quality assessment on a Bioanalyzer Tape Station and confirm pool concentration with qPCR. Samples were then sequenced with Illumina MiSeq using the 600-cycle kit run as 250x250 paired-end sequencing.

For the first batch of samples, 16S and COI libraries were independently pooled and each pool was sequenced in a single lane of Illumina sequencing. The second batch included a smaller number of samples, so to achieve a similar depth of sequencing as the first batch, we pooled and sequenced 16S libraries and COI libraries together in the same sequencing lane.

### *16S* amplicon sequence processing

We used *QIIME 2* v2020.8 (Bolyen et al. 2019) to process 16S sequencing reads and obtain a table of counts of amplicon sequence variants (ASVs, or amplicon sequences representing microbial taxonomic units) across samples. For each sequencing run, we imported demultiplexed paired-end sequences, used the function *qiime dada2 denoise-paired* to trim primer sequences from the 3’ ends of reads, and to trim five bases from the 5’ ends of reads before merging read pairs and detecting ASVs. We then assigned taxonomic classification to ASVs using the SILVA database (v138 SSURef NR99, Quast et al. 2013).

Upon classification, we removed mitochondrial, chloroplast, unassigned, and eukaryotic ASVs. We also identified and removed possible contaminant ASVs by contrasting the presence/absence of ASVs in our negative controls with their prevalence in positive fecal samples (i.e., non-negative controls) using the R package *decontam* (Davis et al. 2018). We used the “prevalence” method to identify and remove ASVs more prevalent in negative controls than in positive samples using a probability threshold of 0.5. We also manually removed ASVs present in negative controls, but absent in positive samples, as these were also likely contaminants. In total, we removed 87 and 359 contaminant ASVs from the batch 1 and batch 2 datasets, respectively.

At this point, we used *QIIME* 2 to merge the feature table, representative sequences, and taxonomy files from the two separate sequencing runs. We finally generated a phylogenetic tree from the merged set of ASV sequences for downstream diversity analyses. We used *qiime phylogeny align-to-tree-mafft-fasttree* to perform multiple sequence alignment, mask highly variable positions, and first generate an unrooted tree and finally a tree rooted at the midpoint of the longest tip-to-tip distance of the unrooted tree.

Finally, we applied several additional filtering steps to achieve a high-quality representation of warbler gut microbiomes. We excluded individuals from species represented by fewer than 5 individuals in our dataset because we were interested in examining species-differences in gut microbiome structure. Because very low depth and uneven depth of sequencing among samples can affect diversity estimates (Hughes & Hellmann et al. 2005), we next generated a rarefied dataset by randomly downsampling ASVs to a minimum threshold to standardize total read counts across samples. We determined the minimum acceptable ASV count threshold by examining rarefaction curves constructed using the *rarecurve* function in *vegan* (Oksanen et al. 2020) using a step size of 50. Based on this analysis, we determined a library size of 4,000 reads to be an acceptable threshold since the number of observed ASVs appears to plateau beyond this point (Figure S2a).

Because we detected a significant effect of sequencing batch on our diversity estimates (i.e., a “batch effect”, see Results), we also performed analyses on a subset of the data that only included the first batch of samples (collected between 2017-2019, referred to as “batch 1”). For these analyses, we performed the same sequence processing steps as above except for merging-in data from the samples collected in 2020.

### COI amplicon sequence processing

We used the AMPtk (v1.5.3) pipeline to analyze COI metabarcoding data by applying the default clustering algorithm (VESEARCH v2.17.1) for operational taxonomic units (OTUs) and assigned taxonomy by pulling from the chordates and arthropods in the BOLDv4 database. We rooted the OTU phylogeny output from AMPtk on a randomly chosen arachnid OTU, as arachnids split from the common arthropod ancestor prior to insects. We then imported the COI metabarcoding data into phyloseq for downstream analyses and applied a similar framework as we did with our 16S data. We first removed OTUs assigned to phylum Chordata as this represents off-target amplification, then rarefied depth to 15,000 reads per individual (full dataset), and 8,500 reads per individual (batch 1 subset) (Figure S2b).

For analyses where we directly investigated the effect of diet on the microbiome at the individual level, we only analyzed individuals with data that passed filtering steps in both microbiome and diet datasets. This included 216 individuals in the full dataset representing 15 species (mean 14 individuals per species) and 130 individuals in the batch 1 subset representing 14 species (mean 9 individuals per species).

### Diet diversity and its relationship with gut microbiome diversity

We estimated within-individual diversity (alpha diversity) of the diet and gut microbiome using the Shannon index and the Chao1 index using the *diversity* function in *vegan*, and using Faith’s phylogenetic diversity using the *estimate_pd* function in *btools* (Battaglia 2022). The Shannon index quantifies ASV richness (the number of ASVs) as well as evenness (the equity in ASV abundances), while Chao1 just quantifies ASV richness. Faith’s phylogenetic diversity is a measure of ASV richness that is the sum of branch lengths in the phylogeny that connect all ASVs in the community assemblage. We estimated between-individual differences between microbiomes (beta diversity) using four different metrics: Bray-Curtis, Jaccard, UniFrac, and weighted UniFrac, calculated using the *distance* function in *phyloseq* (McMurdie & Holmes 2013). Bray-Curtis measures differences in community composition and is based on ASV abundances, whereas Jaccard is based only on presence/absence and does not rely on abundance. UniFrac measures the phylogenetic distance between communities based on presence/absence of ASVs, whereas weighted UniFrac is similar but weights branch lengths by ASV abundance.

We used three approaches to examine the relationship between diet and the gut microbiome. With the prediction that a generalized diet, characterized by a high diversity of arthropod taxa, supports a high gut microbiome diversity, we first tested for a positive correlation between individual diet alpha diversity and gut microbiome alpha diversity using a Kendall’s rank correlation test.

Second, at the species level, we tested whether gut microbiome structure differs among species with a more specialized and less diverse diet, and species with a more generalized and more diverse diet using permutational multivariate analysis of variance (PERMANOVA) of beta diversity distances using the *adonis2* function in *vegan*. For this analysis, we categorized each species as either “low diversity” diet, “high diversity” diet, or “intermediate” by creating an index of diet specialization (Figure 1a). To calculate this index, we summed mean individual within-species diet alpha diversity and mean within-species diet beta diversity with the assumption that (1) more specialized diets are characterized by a lesser diversity of food items (low alpha diversity) and individuals within more specialized species eat a similar diet (low beta diversity), and (2) more generalized diets are characterized by a high diversity of food items (high alpha diversity), and individuals within more generalized species may have highly divergent diets depending on local food availability (high beta diversity). Thus, a low score reflects a less diverse and more specialized diet, and a high score reflects a more diverse and more generalized diet. We note this index quantifies diversity of the diet and that host species within the same diet categorization may have dissimilar diets by way of diet content (e.g., proportion that is flying insects).

For both alpha diversity and beta diversity of the diet, the different diversity metrics we calculated were positively correlated (with the exception of weighted UniFrac and UniFrac beta distance when using the full dataset; Table S1) and diet type classification of each species was consistent across metrics. Thus, for simplicity we report the index of diet specialization using the Shannon index to estimate alpha diversity and the Bray-Curtis metric to estimate beta diversity.

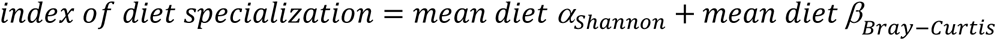

We note that because we used a subset of individuals to calculate diet index for batch 1, for some species classification of diet diversity using the diet index is not consistent between this subset and the full dataset. Four species are classified as intermediate in one dataset and either as high diversity or as low diversity in the other dataset. However, species diet index values are positively correlated between the full dataset and batch 1 (τ=0.516, *P*=0.010, Figure S3), suggesting this index is robust to individual variation in diet. Our results do not change when excluding these four species from the analyses so we include them in our results.

Finally, we used topological congruence analysis to determine whether similarity in gut microbiome structure among host species reflects diet similarity with the expectation that if diet directly shapes host gut microbiomes, then clustering of species by diet similarity will mirror clustering of species by gut microbiome similarity. To generate dendrograms representative of each species, we generated a new ASV table—each for rarefied COI sequence counts and rarefied 16S sequence counts—grouped by host species by averaging ASV counts within each species, re-calculated dissimilarity matrices and constructed dendrograms by clustering distance matrices using the UPGMA method in the *hclust* function in R (following Trevelline et al. 2020).

We then compared the observed 16S dendrogram to the observed COI dendrogram using *TreeCmp* (Bogdanowicz et al. 2012) to compute the matching cluster metric of topological congruence (Bogdanowicz & Giaro 2013). Following Brooks et al. (2016), we then compared the observed 16S dendrogram with 10,000 dendrograms with randomized topology and calculated a normalized congruence score, which is the observed matching cluster score divided by the maximum congruence score between the observed dendrogram and one of the random dendrograms. Finally, we evaluated significance and report a p-value by dividing the number of randomized dendrograms with equal or more congruent scores to the observed 16S dendrogram than the score between the two observed dendrograms by 10,000. We also used Mantel tests as a complimentary analysis to examine correlations between the diet and microbiome beta distance matrices at the individual level, where each value represents the beta distance between a pair of individuals, using *vegan::mantel* with the spearman correlation method.

### Gut microbiome diversity and topological analyses

We identified a “core” wood-warbler gut microbiome as the collection of ASVs present across a large number of individuals using the rarefied dataset. Because most ASVs had a low prevalence among individuals (Figure S4), we report the core microbiome as ASVs present in >30% of all individuals. Although this threshold is arbitrary, we believe it is conservative as only 39 ASVs were represented in greater than 30% of individuals (see below). We also report taxa at high relative abundance across all samples at phylum level. This set of ASVs represents bacteria that are most common in the gut microbiome among breeding male wood-warblers.

To quantify the effect of host taxonomy on the gut microbiome and the extent to which gut microbiomes covary with host phylogeny, we took two approaches using the full set of ASVs. First, we estimated gut microbiome divergence (beta diversity) among individuals using four measures of community dissimilarity: Bray Curtis distance, Jaccard distance, and weighted and unweighted UniFrac distances. We then used *vegan::adonis2* to conduct PERMANOVA tests to determine the effect of host species on community dissimilarity. Because our samples were collected across four breeding seasons, from two geographic localities, and were sequenced in two different batches we also tested for effects of these factors. We included each of these factors in our model and set the “by” parameter to “margin”. However, in the full dataset the effects of sampling year and sequencing batch are confounded since all samples collected in 2020 were sequenced in batch 2, so we ran two separate models which included either host species + locality + year, or host species + locality + sequencing run. Results for host species and locality were similar between models, so we report results from the model that included sequencing run for simplicity. We also calculated multivariate homogeneity of group dispersions for significant variables using *vegan::betadisper* and assessed deviations from this expectation using *vegan::permutest* because a homogeneous dispersion among groups is an assumption for PERMANOVA tests. We visualized beta distances between gut microbiota using the principal coordinate analysis (PCoA) method of *phyloseq::ordinate*.

Our second approach was to test for congruence between the host phylogeny and microbiome, as phylosymbiosis predicts host relatedness and microbiome community similarity to exhibit a positive relationship (Brooks et al. 2016). To do this, we first used the same topological congruence approach as described above, but used the topology from and Baiz et al. (2021) for *Setophaga* species, and from Lovette et al. (2010) for outgroup taxa in place of the diet dendrogram (Figure 1a). We then also used Mantel tests to test for correlations between the gut microbiome distance matrix and a matrix of cophenetic distances, representing evolutionary distances, between individuals. We calculated cophenetic distance between species using the *stats::cophenetic* function on a dendrogram representing the host phylogeny in Figure 1a, with branch lengths scaled using divergence times from TimeTree of Life (Kumar et al. 2017; Table S2). Note that an evolutionary distance of zero denotes a pair of individuals from the same species.

Because we found a significant influence of sequencing batch on gut microbiome diversity, we separately performed all analyses on the subset of samples sequenced in the first batch (collected between 2017-2019, referred to as “batch 1”) as this batch included a larger subset of samples that were collected across multiple years than the second batch, which only included samples collected in 2020. For topology and Mantel analyses, we also subset our data to account for potentially confounding effects of (1) geographic locality by only analyzing samples collected in New York between 2017-2019 (referred to as “batch 1-NY”) and (2) sampling year by only analyzing samples collected in 2020 (referred to as “batch 2”).

## Results

### *16S* sequencing output and composition of the warbler gut microbiome

The number of ASVs yielded by our first 16S sequencing run was 6,412 (per-individual median=36, mean=53, SD=65) while our second 16S sequencing run yielded 10,590 ASVs (per-individual median=235, mean=218, SD=73). This discrepancy is likely explained by a higher average depth of sequencing across individuals in the second sequencing run (Figure S5), despite our attempt at normalization. Taxa that were detected in both sequencing runs represented a small proportion of the total number of ASVs across runs (6%), contributing to the gut microbiome differentiation we observed for individuals sampled in 2020 (see below).

After merging our 16S datasets, applying our filtering steps and rarefaction, our full dataset consists of 270 individuals representing 15 species (mean 18 individuals per species, with 95% of individuals being male, 1% female, and 4% of unknown sex). Among these samples, we detected 12,048 ASVs from 39 bacterial phyla with the top phylum, Proteobacteria, representing 60% of the total reads (Figure 1b). Firmicutes was the next most abundant phylum, representing 13% of the total reads, followed by Actinobacteriota, representing 6.5% of the total reads. The remaining phyla each represented <5% of the total reads. We observed considerable variation in relative abundance of prevalent taxa between individuals of the same species (Figure S6a). Despite low overlap in ASV identity between sequencing runs, composition and relative abundance of prevalent phyla were very similar across host species when we separately examined samples that were sequenced in different batches (Figure S7).

Most ASVs were present in <10% of individuals, and only 39 ASVs were represented in >30% of individuals (Figure S4). Each of these core ASVs was represented in all but one or two of the host species we analyzed (Table S3 and S4). The most prevalent ASV was a Gammaproteobacteria of the family Yersiniaceae. This ASV was found in all 15 host species and ∼60% of samples in both the full dataset and the batch 1 subset. Gut microbiome alpha diversity did not differ among host species (Kruskal-Wallis rank sum test: Shannon index: full dataset d.f.=14, *χ*^2^=14.68, *P*=0.400; batch 1 d.f.=13, *χ*^2^ =16.354, *P*=0.231, Chao1 index: full dataset d.f.= 14, *χ*^2^= 13.99, *P*=0.451; batch 1 d.f =13, *χ*^2^ =19.32, *P*=0.113, Faith’s PD: full dataset d.f.=14, *χ*^2^= 14.98, *P*=0.380; batch 1 d.f.=13, *χ*^2^=18.764, *P*= 0.131).

### COI sequencing output, diet diversity and its relationship with gut microbiome diversity

Our first COI sequencing run yielded 3,235 OTUs, while the second yielded 2,668 OTUs. In contrast to the 16S dataset, there was moderate overlap in OTU identity between sequencing runs (37% of OTUs are represented in both batches).

Our analyses revealed 4,397 OTUs in the full COI dataset, which was reduced to 3,227 after filtering and rarefaction. Among warbler species, ∼70% or greater relative abundance of diet taxa consisted of insects, particularly in the orders Diptera and Lepidoptera (Figure 1c, Figure S6b).

The majority of other diet taxa included Arachnids in the family Araneae. There was a high degree of overlap among species in diet PCoA space (Figure 3b). These results were consistent between analyses that included all individuals and only individuals sequenced in the first batch.

Warbler species fell into three natural partitions along our index of diet specialization, thus we used these partitions to classify species according to diet type (Figure 2b). We classified 2-3 warbler species with low diversity diets depending on the dataset being analyzed (batch 1: American Redstart (AMRE), Chestnut-sided Warbler (CSWA); full dataset: American Redstart (AMRE), Chestnut-sided Warbler (CSWA), Worm-eating Warbler (WEWA), 2-4 species with high diversity diets (batch 1: Black-throated Green Warbler (BTNW), Canada Warbler (CAWA); full dataset: Black-and-white Warbler (BAWW), Canada Warbler (CAWA), Common Yellowthroat (COYE), Hooded Warbler (HOWA)), and the remainder of species as intermediate (Figure 1a).

**Figure 2.**
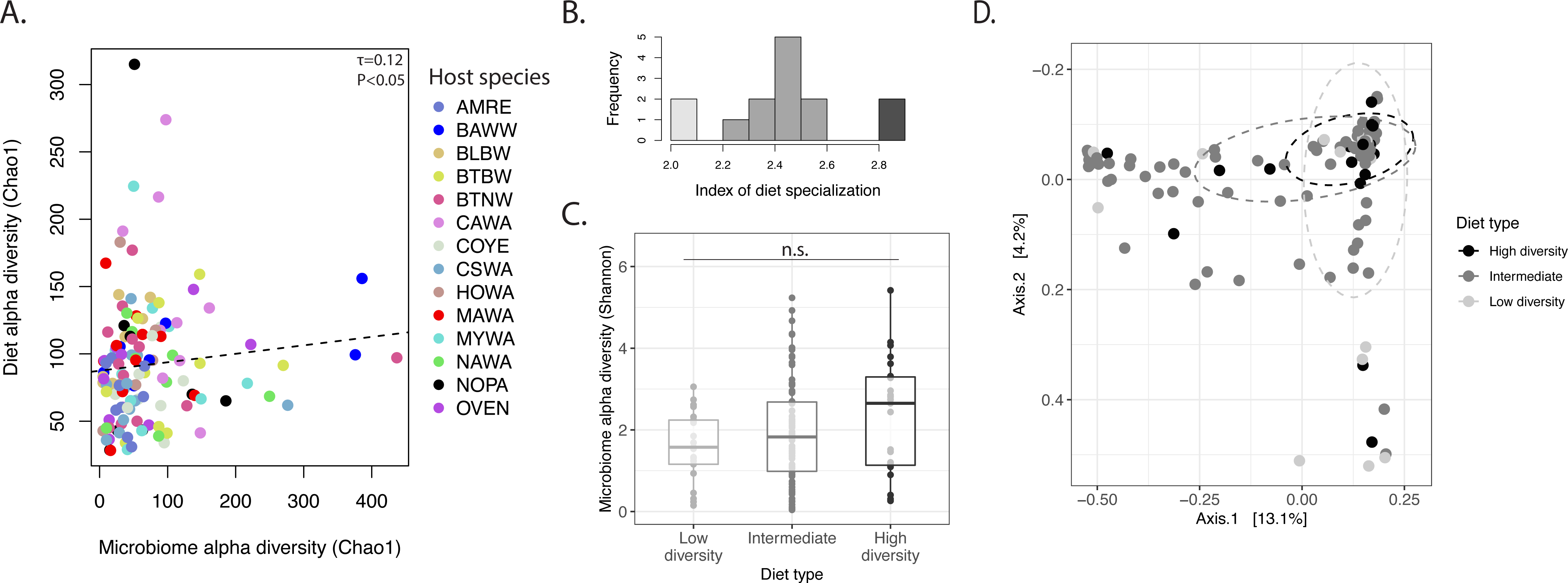
Relationship between diet diversity and gut microbiome diversity. **A)** Within-individual diversity of the gut microbiome is weakly correlated with within-individual diet diversity as measured by the Chao1 index in the batch 1 dataset. Dashed line is a linear model fit to the data. Point color reflects warbler species (see Table 1 for species codes). **B)** Distribution of diet index scores by host species, where a low score is reflective of low diet diversity or diet specialization. Color indicates assignment to diet type and is consistent with part C and D, **C)** Microbiome alpha diversity does not differ among diet types, as classified by diet index. **D)** Principal coordinate analysis (PCoA) of Bray-Curtis distance between host gut microbiomes sequenced in batch 1. Point color represents species diet type as defined by our index of diet specialization. Ellipses are drawn at 50% confidence level. In each panel, data shown are from sequencing batch 1.

**Figure 3.**
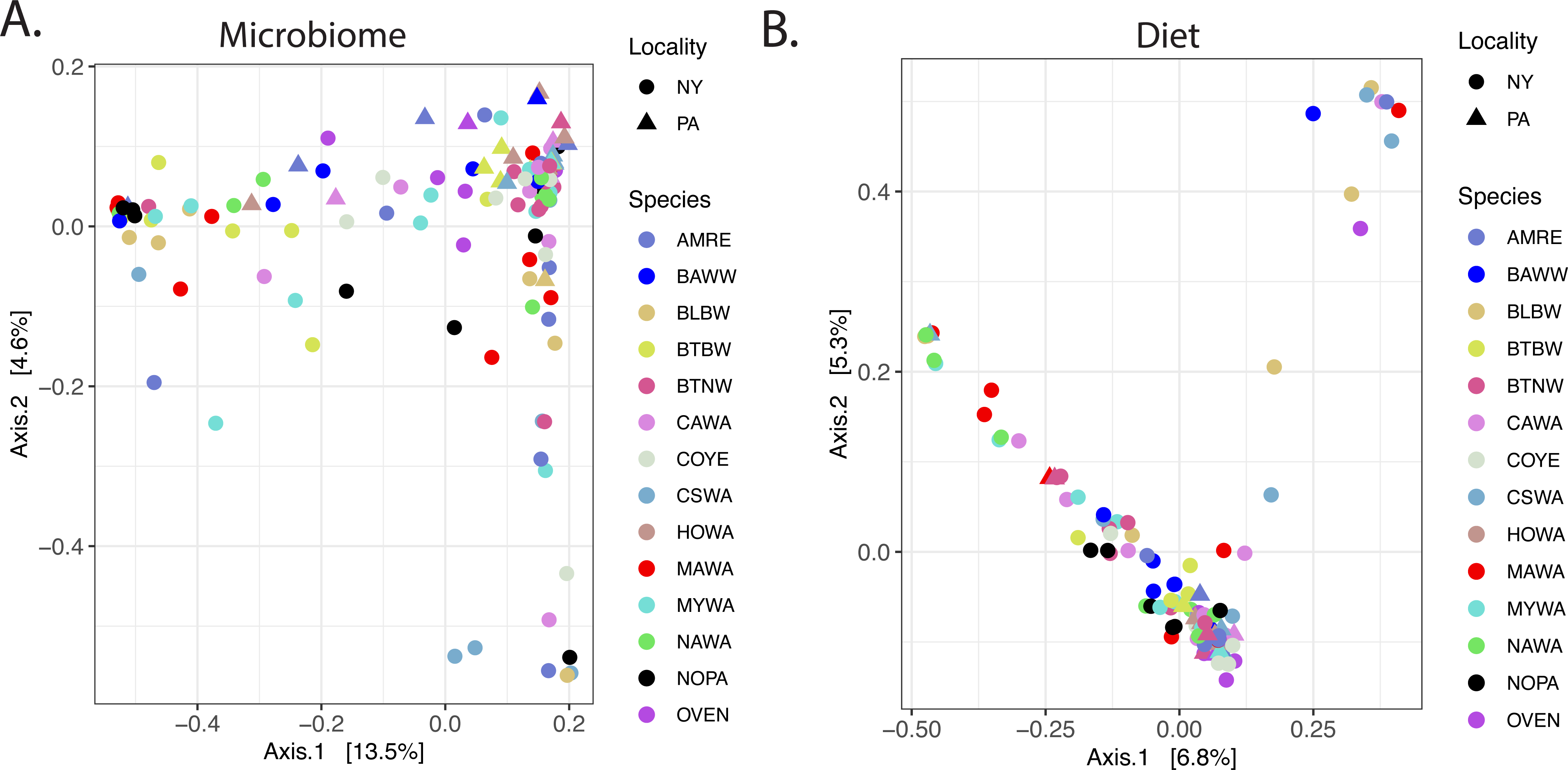
Principal coordinate analysis (PCoA) of Bray-Curtis distance between **A)** host gut microbiomes and **B)** individual diet. In both panels, data are from samples sequenced in batch 1. Point color represents host species, and shape represents geographic locality. See Table 1 for species codes.

When considering within-individual diversity, we found no correlation between diet alpha diversity and microbiome alpha diversity when using Shannon index and Faith’s PD, as well as Chao1 when considering the full dataset (Kendall’s rank correlation, Shannon index batch 1: τ=0.029, *P*=0.619; full dataset: τ=0.005, *P*=0.906, Faith’s PD batch 1: τ=0.110, *P*=0.064; full dataset: τ=-0.034, *P*=0.459, Chao1 full dataset: τ=-0.063, *P*=0.171,), but when considering the batch 1 subset using Chao1, alpha diversity of the diet and microbiome were positively correlated (Figure 2a; Kendall’s rank correlation batch 1: τ=0.124, *P*=0.038). This indicates that for batch 1, individuals that consumed high richness diets (more OTUs) tended to have more rich gut microbiota (more ASVs), but the correlation is weak. Alpha diversity of the microbiome was generally lower for individuals of species that were diet specialists, and higher for individuals of species that were diet generalists (Figure 2c), but alpha diversity of the microbiome did not significantly differ by species diet type (Kruskall-Wallis d.f.=2: Shannon index batch 1: *χ*^2^=2.8, *P*=0.242; full dataset: *χ*^2^=0.014, *P*=0.993, Chao1 batch 1: *χ*^2^=5.4, *P*=0.068; full dataset: *χ*^2^=0.31, *P*=0.855, Faiths PD batch 1: *χ*^2^=4.4, *P*=0.110; full dataset: *χ*^2^=0.13, *P*=0.936), even when only comparing low diversity diets to high diversity diets (Kruskall-Wallis d.f.=1: Shannon index batch 1: *χ*^2^=1.934, *P*=0.1643, full dataset: *χ*^2^=0.049, *P*=0.825, Chao1 batch 1: *χ*^2^=1.8, *P*=0.181; full dataset: *χ*^2^=0.29, *P*=0.592, Faith’s PD batch 1: *χ*^2^=1.3, *P*=0.259; full dataset: *χ*^2^=0.17, *P*=0.676).

### Factors accounting for warbler gut microbiome structure

When analyzing the full dataset which included microbiomes sequenced in two different sequencing runs, there was a very clear and strong batch effect where microbiomes sequenced in one run were more similar to each other than to microbiomes sequenced in the other run (Figure S8). Yet, principal coordinates analysis of gut microbiome dissimilarity matrices revealed a high degree of overlap among hosts of different species and among hosts from different geographic localities (Figure 3a). There was little clustering of microbiomes by diet type of host species as defined by our index of diet specialization (Figure 2d).

Our PERMANOVA tests (Table 2) revealed that sequencing run explained a relatively high degree of variation in Bray-Curtis distances (13%, *P*=0.001), Jaccard distances (7.1%, *P*=0.001), and UniFrac distances (7.9%, *P*=0.001). This strong batch effect likely confounded tests of other variables, since the second sequencing run only contained samples collected in a single year (2020) and included an additional species (WEWA, Worm-eating warbler) that is not represented in the first sequencing run. Thus, we analyzed the subset of data from 2017-2019 (i.e., batch 1) separately to examine the effect of biological factors on microbiome structure in the absence of the sequencing batch effect, because of the two sequencing runs this batch included the largest sample size of individuals and included three years of sampling. This analysis revealed sampling locality had a significant effect when using all four distance metrics, although the effect size was small (∼1-2% of variation explained; Table 2). Similarly, year explained a small amount of variation (∼1.5%) when using Jaccard and UniFrac distances. In the absence of the sequencing batch effect, host species identity accounts for the highest degree of variation in microbiome structure when using Bray-Curtis (9%, *P*=0.048), Jaccard (9.3%, *P*=0.001) and UniFrac distances (10.3%, *P*=0.001), generally explaining ∼6-fold more of the variation than any other factor considered. Permutation tests indicated that dispersion among species Jaccard and UniFrac distances is not homogenous, which could account for the significant PERMANOVA result. However, this does not seem to be the case because although dispersion is high for several species causing overlap in PCoA space, species’ centroid positions are largely non-overlapping when using Bray-Curtis, Jaccard and UniFrac distances (Figure S9), likely reflecting true gut microbiome structuring among species.

**Table 2.** Results of permutational multivariate analysis of variance (PERMANOVA) tests and permutation tests of dispersion on beta distances between gut microbiomes. Diet type reflects categorization based on our index of diet specialization (i.e., high diversity, low diversity, intermediate). Asterisks denote significant results: *** *P*<0.001, ** *P*<0.01, \**P*<0.05 level.

|  |  | PERMANOVA |  |  | Permutation test on dispersion |  |
| --- | --- | --- | --- | --- | --- | --- |
| Distance matrix | Variable | d.f. | R<sup>2</sup> | P | F | P |
| Full dataset (2017-2020) |  |  |  |  |  |  |
| Bray-Curtis | Species | 14 | 0.047 | 0.162 |  |  |
|  | Locality | 1 | 0.007 | 0.011* | 13.186 | 0.001** |
|  | Year | 3 | 0.138 | 0.001** | 52.964 | 0.001** |
|  | Sequencing run | 1 | 0.130 | 0.001** | 171.570 | 0.001** |
|  | Diet type | 2 | 0.009 | 0.435 |  |  |
| Jaccard | Species |  | 0.053 | 0.001** | 4.481 | 0.001** |
|  | Locality |  | 0.005 | 0.009** | 30.959 | 0.001** |
|  | Year |  | 0.080 | 0.001** | 151.68 | 0.001** |
|  | Sequencing run |  | 0.071 | 0.001** | 527.04 | 0.001** |
|  | Diet type |  | 0.010 | 0.167 |  |  |
| UniFrac | Species | 14 | 0.057 | 0.001** | 3.189 | 0.001** |
|  | Locality | 1 | 0.006 | 0.012* | 0.098 | 0.761 |
|  | Year | 3 | 0.087 | 0.001** | 8.033 | 0.001** |
|  | Sequencing run | 1 | 0.079 | 0.001** | 21.312 | 0.001** |
|  | Diet type | 2 | 0.010 | 0.191 |  |  |
| Weighted UniFrac | Species | 14 | 0.062 | 0.163 |  |  |
|  | Locality | 1 | 0.013 | 0.020* | 2.155 | 0.146 |
|  | Year | 3 | 0.016 | 0.146 |  |  |
|  | Sequencing run | 1 | 0.006 | 0.194 |  |  |
|  | Diet type | 2 | 0.013 | 0.190 |  |  |
| Batch 1 (2017-2019) |  |  |  |  |  |  |
| Bray-Curtis | Species | 13 | 0.090 | 0.048* | 0.703 | 0.762 |
|  | Locality | 1 | 0.014 | 0.002** | 0.759 | 0.397 |
|  | Year | 2 | 0.012 | 0.385 |  |  |
|  | Diet type | 2 | 0.020 | 0.062 |  |  |
| Jaccard | Species |  | 0.093 | 0.001** | 2.409 | 0.009** |
|  | Locality |  | 0.011 | 0.001** | 5.607 | 0.020* |
|  | Year |  | 0.015 | 0.001** | 5.889 | 0.004** |
|  | Diet type |  | 0.020 | 0.001** | 9.067 | 0.001** |
| UniFrac | Species | 13 | 0.103 | 0.001** | 3.049 | 0.002** |
|  | Locality | 1 | 0.009 | 0.014* | 0.002 | 0.971 |
|  | Year | 2 | 0.015 | 0.034* | 2.212 | 0.099 |
|  | Diet type | 2 | 0.022 | 0.013* | 2.161 | 0.118 |
| Weighted UniFrac | Species | 13 | 0.085 | 0.380 |  |  |
|  | Locality | 1 | 0.017 | 0.039* | 0.183 | 0.655 |
|  | Year | 2 | 0.008 | 0.732 |  |  |
|  | Diet type | 2 | 0.021 | 0.246 |  |  |

In line with our findings of little-to-no correlation between individual diet diversity and gut microbiome diversity, host species diet type did not significantly explain variation between microbiomes in the full dataset, nor in the batch 1 subset--with the exception of using Jaccard and UniFrac distance, in which case diet type explained a small amount of variation (∼2%; Table 2). Dispersion among diet types for Jaccard distance was not homogenous (F=9.067, *P*=0.001), yet diet type centroid positions for Jaccard and UniFrac distances were non-overlapping in PCoA space especially for low diversity diets (Figure S10), indicating some differentiation among gut microbiota for species with more specialized diets.

### Topological congruence analyses

Normalized matching cluster congruence scores for the gut microbiome-host phylogeny topological comparisons were between ∼0.4-0.8. As congruence scores of zero indicate complete topological congruence, and scores of 1 indicate complete incongruence, these scores reflect intermediate congruences. When analyzing all individuals in the full dataset, and within the batch 1 and batch 1-NY subsets, the observed warbler gut microbiome dendrogram was significantly more congruent with the host phylogeny than with randomized dendrograms using Bray-Curtis, Jaccard and weighted UniFrac distances (Table 3, Figure 4a). In the batch 2 subset, the gut microbiome dendrogram was more congruent with the host phylogeny than with randomized dendrograms using Bray-Curtis and UniFrac distances (Table 3). Thus, the majority of comparisons (N=11 of 16 comparisons) indicate a positive association between gut microbiome similarity and host phylogenetic relatedness. As Bray-Curtis and weighted UniFrac metrics are weighted by ASV counts, this may indicate that relative abundances of microbial taxa help contribute to the phylogenetic signal in the warbler gut microbiome.

**Figure 4.**
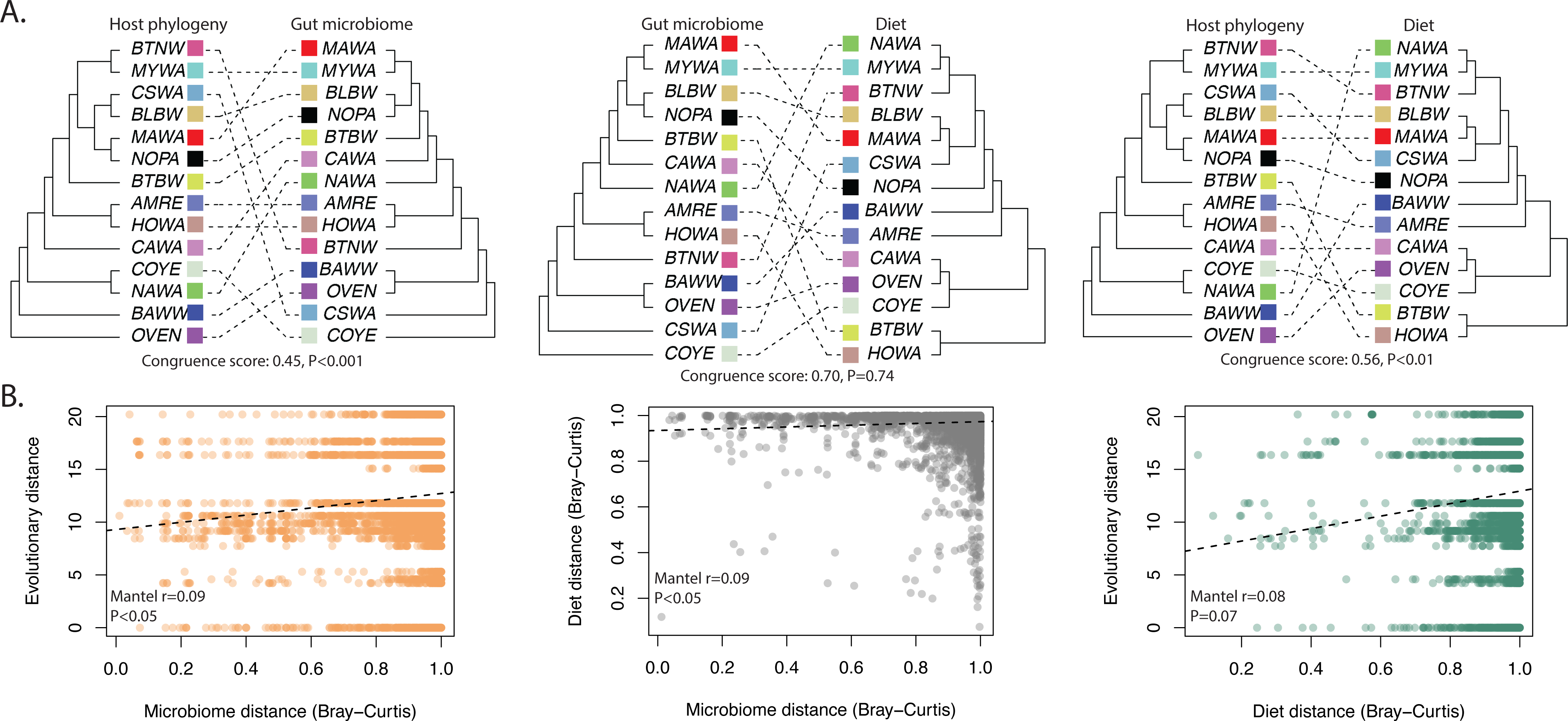
Summary of phylosymbiosis analyses. **A)** Topological congruence analyses of the association between the gut microbiome and host phylogeny (left), the gut microbiome and diet (middle), and the diet and host phylogeny (right). Microbiome and diet dendrograms were constructed using Bray-Curtis distances of mean within-species ASV/OTU counts. Matching cluster congruence scores are normalized where 0=complete congruence, and 1=complete incongruence. See Table 1 for species codes. **B)** Scatter plots of individual-level microbiome versus host evolutionary distances (left), microbiome versus diet distances (middle), and diet versus host evolutionary distances. Diet and microbiome distances are of the Bray-Curtis metric.

**Table 3.** Summary of topological congruences between species-level gut microbiome dendrograms and the host phylogeny (left), and between species-level diet dendrograms (right), and of individual-level Mantel tests. N spp.=number of species analyzed, and matching cluster congruence scores are normalized where 0=complete congruence, and 1=complete incongruence. Asterisks denote significant results: *** *P*<0.001, ** *P*<0.01, \**P*<0.05 level.

|  |  |  | Microbiome-host phylogeny |  | Microbiome-diet |  |
| --- | --- | --- | --- | --- | --- | --- |
|  | N spp. | Distance metric | Matching cluster congruence score | Mantel r | Matching cluster congruence scores | Mantel r |
| Full dataset (2017-2020) | 15 | Bray-Curtis | 0.52*** | 0.02 | 0.57 | 0.06* |
|  |  | Jaccard | 0.58** | 0.04 | 0.56*** | 0.17** |
|  |  | UniFrac | 0.68 | 0.10** | 0.74 | 0.16** |
|  |  | Weighted UniFrac | 0.58** | 0.01 | 0.51** | 0.03 |
| Batch 1 (2017-2019) | 14 | Bray-Curtis | 0.45*** | 0.09* | 0.70 | 0.09* |
|  |  | Jaccard | 0.45*** | 0.18** | 0.39*** | 0.21** |
|  |  | UniFrac | 0.71 | 0.19** | 0.64 | 0.22** |
|  |  | Weighted UniFrac | 0.57** | -0.003 | 0.61 | -0.05 |
| Batch 1-NY (2017-2019) | 13 | Bray-Curtis | 0.44*** | 0.15** | 0.53* | 0.11* |
|  |  | Jaccard | 0.52** | 0.23** | 0.44*** | 0.23** |
|  |  | UniFrac | 0.73 | 0.27** | 0.69 | 0.25** |
|  |  | Weighted UniFrac | 0.52** | 0.07 | 0.74 | -0.09 |
| Batch 2 (2020) | 14 | Bray-Curtis | 0.56** | 0.07 | 0.62* | 0.01 |
|  |  | Jaccard | 0.72 | 0.03 | 0.67 | -0.03 |
|  |  | UniFrac | 0.59* | 0.04 | 0.64 | -0.02 |
|  |  | Weighted UniFrac | 0.79 | -0.02 | 0.73 | 0.06 |

To determine whether gut microbiome similarity better reflects host evolutionary history or host diet similarity, we repeated the topological analyses above instead using a dendrogram clustered from the diet OTU distances in place of the host phylogeny (Table 3). Among comparisons, congruence scores were generally lower (indicating better congruence) for microbiome-host phylogeny comparisons than for microbiome-diet comparisons (Figure S11), although there are some exceptions. Importantly, only six of sixteen microbiome-diet comparisons were significantly more congruent than random. Three of these comparisons were of Jaccard distance, which only considerers ASV presence/absence. Further, in two other instances both considering Bray-Curtis distances, congruence scores for the microbiome-host phylogeny comparison were lower than for the microbiome-diet comparison (batch 2, batch 1-NY; Table 3). Collectively, these results suggest a closer association between gut microbiome structure and host evolutionary history than with host diet.

Finally, we examined the association between the host phylogeny and diet dendrograms from the batch 1 subset to determine whether the significant associations we detected between the gut microbiome and diet could be due to a phylogenetic signal of the diet. For all four distance metrics, the diet-host phylogeny comparison was significantly more congruent than random (Bray-Curtis normalized matching cluster score=0.56, *P*=0.007, Jaccard normalized matching cluster score=0.52, *P*=0.003, UniFrac normalized matching cluster score=0.51, *P*=0.001, weighted UniFrac normalized matching cluster score=0.57, *P*=0.017). These scores reflect intermediate congruence between the diet dendrogram and host phylogeny, and are similar but slightly higher (less congruent) on average than congruence scores between the microbiome dendrogram and host phylogeny (Figure 4A).

### Mantel tests

Mantel tests indicated a positive relationship between individual-level microbiome distances and pairwise evolutionary distances (Mantel r ∼0.09-0.27) in the batch 1 and batch1-NY datasets for Bray-Curtis, Jaccard, and UniFrac distances, and in the full dataset for UniFrac distance (Table 3, Figure 4b). Mantel tests also indicated a positive relationship between individual-level microbiome distances and diet distances (Mantel r ∼0.06-0.25) in the full dataset, batch 1, and batch 1-NY subsets using Bray-Curtis, Jaccard, and UniFrac distances (Table 3, Figure 4b).

We also tested the relationship between diet matrices and pairwise evolutionary distances in the batch 1 subset, and found a positive association for Jaccard diet distance (Mantel r=0.10, *P*=0.027), and UniFrac diet distance (Mantel r=0.13, *P*=0.004). Notably, across all Mantel tests, most significant correlations were detected when using unweighted distance matrices (Jaccard and UniFrac).

## Discussion

We performed fecal metabarcoding to examine environmental and evolutionary influences on gut microbiome structure in breeding wood warblers. Our analyses collectively support host taxonomy as the strongest driver of gut microbiome structure while environmental factors, including diet diversity, showed lesser effects. At the individual level, diet diversity—both within and between individuals—showed little-to-no association with microbiome diversity.

Further, on average, more closely related species tended to harbor more similar gut microbiomes, and gut microbiome similarity was less closely associated with diet similarity, suggesting host evolutionary history may play a large role in shaping host-microbe interactions in this clade. We also detected a relatively strong batch effect of sequencing run on gut microbiome diversity, and by analyzing within-batch subsets of our data we saw this had obscured the signal of the biological factors we considered in our analyses. Thus, these results highlight caution for other researchers about whether or not to divide samples across sequencing lanes and this should be a serious consideration in future metabarcoding studies.

### The wood warbler gut microbiome

Wood warbler gut microbiomes were dominated by Proteobacteria and Firmicutes, which is consistent with other studies of other free-living passerines (e.g., Hird et al. 2015, Bodawatta et al. 2021a). The most prevalent ASV, a Proteobacteria in the family Yersiniaceae, was observed in ∼60% of individuals and occurred in all host species examined, but only a very small proportion of ASVs were represented in >30% of the individuals sequenced. These results may reflect a shared signature of the passerine gut microbiome in wood warblers at higher taxonomic levels, yet a high level of variability among individuals, especially for lower abundance taxa.

The most dominant bacterial phyla in the current study were also identified as highly abundant in the only migratory cycle study of re-captured warblers to-date, which focused on Kirtland’s warblers (*Setophaga kirtlandii*; Skeen et al. 2021), a species that does not breed in our study areas. Although arrival on the breeding grounds was accompanied by a shift from a Kirtland’s warbler gut microbiome dominated by Firmicutes to one dominated by Proteobacteria, both phyla were highly abundant across the migratory cycle. The most prevalent taxonomic classes in the current study (Gammaproteobacteria, Alphaproteobacteria, and Bacilli) also dominated gut microbiomes of breeding Kirtland’s warblers (Skeen et al. 2021). However, Clostridia was one of the most abundant taxa in Kirtland’s warblers but was found at low prevalence among individuals in the current study and made up only <2% of the total reads sequenced. This may suggest that Kirtland’s warblers, a near threatened Caribbean migrant with highly specialized habitat requirements, differ in gut microbiome structure from closely related parulids breeding nearby. This differentiation would be consistent with our findings of a relatively strong role of host taxonomy and evolutionary history, and/or associated environmental factors that we were unable to resolve with our dataset, in shaping the parulid gut microbiome (see below).

In this study, sampling locality consistently explained 1-2% of variation between microbiomes across datasets and distance metrics considered. Samples were collected from two forested localities in Eastern North America roughly 400 km apart, a distance that is likely not large enough to generate significant population genetic structure within warbler host species due to a lack of potential barriers to gene flow (e.g., yellow-rumped warblers, *S. coronata;* Toews et al. 2016). However, our results suggest this distance may be sufficient in scale to affect subtle changes in gut microbe communities. Interestingly, the amount of variation explained by sampling locality here is similar to that reported in other passerine studies (San Juan et al. 2021, Teyssier et al. 2020), despite this study encompassing a larger geographical area. For example, habitat type explained ∼4% of variation between passerine microbiomes within a 43 km agricultural study area in Costa Rica (San Juan et al. 2021), suggesting habitat features may be more important than geographic distance between sites. Although we did not include habitat features as a factor in our analyses, notable differences between our study sites include an abundance of *Rhododenron* (*R. maximum*) and mountain laurel (*Kalmia latifolia*) in the understory at our Pennsylvania localities, whereas these shrubs do not occur in our New York localities. This and other habitat differences could conceivably contribute to the differences we observed in gut microbiota between our sites.

When analyzing a subset of samples from a single sequencing run, sampling year explained a similar proportion of variation between microbiomes as did sampling locality, but tended not to be significant. This may indicate that wood warbler microbiomes are stable across breeding seasons, despite annual long-distance longitudinal migration to-and-from tropical non-breeding grounds, which is likely associated with changes in foraging strategies. This is consistent with other passerine studies which found no difference in gut microbiome diversity across consecutive breeding seasons (Escallón et al. 2019, Benskin et al. 2015), but it is important to note that in our dataset, each year represents a different cohort of individuals. In migratory species, it will be desirable to re-sample the same individuals on the non-breeding and breeding grounds across multiple cycles to disentangle temporal effects from those of habitat, diet and geographic locality (Skeen et al. 2021).

### Diet diversity is not tightly linked to gut microbiome diversity in wood warblers

By sequencing arthropod COI metabarcoding libraries from the same fecal samples we amplified bacterial 16S libraries, we were able to directly examine the relationship between natural diet diversity and gut microbiome diversity. Our strategy revealed that when analyzing three different metrics of within-individual (alpha) diversity, diet diversity was not correlated with microbiome diversity with the exception of a weak correlation in the batch 1 data when using the Chao1 index which is neither phylogenetically aware nor weighted by ASV/OTU abundance (Figure 2a). Although individuals of species with low diversity diets tended to have reduced gut microbiome alpha diversity and individuals of species with high diversity diets tended to have increased microbiome alpha diversity, this pattern was not significant (Figure 2c). Further, when looking at between-individual (beta) diversity, diet type only explained ∼2% of the variation between individuals and only when using unweighted distance metrics. In this case, individuals of species with more specialized (less diverse) diets tended to drive this pattern (Figure S10). This provides some evidence that diet richness may be weakly associated with gut microbiome richness, although we were unable to detect significant associations with these analyses when using our full dataset which may be due to the batch effect. Thus, in contrast to our prediction, diversity of the diet generally did not explain variation in the gut microbiome.

This may suggest a high diversity diet either does not generally provide wood warblers an increased availability of potential gut colonists, or gut microbe colonization is not affected by diet diversity. Similarly, in a study of two species of freshwater fish, Bolnick et al. (2014) found the relationship between diet diversity and gut microbiome diversity was not linear and fish with a specialized diet actually harbored a more diverse gut microbiome.

Despite our finding of little relationship between diet diversity and gut microbiome diversity, many studies have shown host diet indeed influences the avian gut microbiome. Broad categorization of natural feeding guild and diet type explain differences in the gut microbiomes of wild passerines in New Guinea and of zoo and farm birds in China, respectively (Bodawatta et al. 2021a, Xiao et al. 2021). Further, experimental manipulations of passerine diets have been associated with shifts in gut microbiome diversity and composition (Davidson et al. 2020, Tyssier et al. 2020, Knutie 2020, Perkarsky et al. 2021). In the current study, we analyzed natural diets of breeding wood warblers, which are known to primarily eat insects (MacArthur 1958, Birds of the World 2022). Our metabarcoding results indicate a substantial portion of the diet is also Arachnid-based. However, diet alpha diversity did not differ among species and relative proportions of arthropod classes in the diet were similar (Figure 1c). The lack of species with a highly specialized diet (at the scale analyzed here) that were included in this study may make wood-warblers a poor system for untangling the effect of diet diversity on gut microbiome diversity, and future dual diet-microbiome metabarcoding studies could also include birds with clear distinctions in dietary guild for comparison (e.g., extreme diet specialists, aerial insectivores). We note that we did not consider fine-scale spatial partitioning of the feeding niche as an explanatory variable in this study, something wood-warblers are well known for (MacArthur 1958). Further, it is possible that because we examined broad-scale patterns in diet diversity at the OTU level, we were not able to identify components of the diet (e.g., nutritional values of arthropods) that possibly underlie gut microbiome structure. We also note that although wood-warblers are primarily insectivores, some species are known to supplement their diet with fruit, especially in the non-breeding season (Birds of the World 2022). Our study design did not allow us to examine effects of any non-arthropod components of the diet, which may influence gut microbiota. Nevertheless, our results suggest dietary arthropod diversity does not scale directly with gut microbiome diversity in breeding wood-warblers.

### Host evolution as the main driver of wood-warbler gut microbiome structure

Amongst the biological factors considered in this study, host species stands out as the variable that explains the largest amount of variation between microbiomes. Further, species-level 16S dendrograms were generally more concordant with the host phylogeny than with COI dendrograms (Figure 4A, Table 3). We also found the host phylogeny to be concordant with COI diet dendrograms, suggesting the weaker associations we did detect between the diet and gut microbiome may have arisen due to a phylogenetic signal of both the diet (Miller al. in prep) and microbiome. Together with our findings of little environmental influence on the wood warbler microbiome, this may suggest that host evolutionary history rather than differences in species’ ecological niche, is the main driver of microbiome differentiation between wood-warbler species.

Mantel analyses of individual-level matrices revealed a somewhat contrasting pattern, showing support for positive associations between the gut microbiome and evolutionary distance and a similar level of support for a positive relationship between the gut microbiome and diet distance. Similar to the topological congruence analysis, these analyses also showed some support for a relationship between the diet and evolutionary distance. In these analyses, most of the significant associations involving the diet arose using unweighted distance metrics. These results are consistent with our other diet diversity analyses by suggesting community richness is driving these patterns.

The conflicting pattern revealed by the topological congruence analyses and Mantel tests may be explained by at least two factors. First, although they are complimentary tests of phylosymbiosis, topological congruence analyses and Mantel tests fundamentally rely on different information. Topological congruence analyses do not rely on branch lengths or directly consider evolutionary or beta distances, whereas Mantel tests measure the correlation between two distance matrices. Because changes in microbiome community structure may be much more rapid than evolutionary changes between host genomes, topological congruence analyses may be a more conservative test of phylosymbiosis (Lim & Bordenstein 2020).

Second, we used species-averaged ASV/OTU counts in the topological congruence analyses in order to summarize variation within each species, whereas our Mantel tests were of distance matrices based on individual-level data. Across all of our analyses of individual-level data, both alpha and beta diversity of the diet and microbiome were quite variable, even within species. For example, Bray-Curtis distances between individuals of the same species ranged from ∼0.07-1 for the gut microbiome, and from ∼0.25-1 for the diet (Figure 4b). This may suggest that the high level of variation within-species obscured phylogenetic signal in gut microbiome and diet similarities at the individual level in Mantel tests.

Host species identity was the biological factor that explained the highest degree of variation between microbiota (Table 2), suggesting the mean ASV counts used in topological congruence analyses may capture unique features within host species. Collectively, our analyses support a tighter association between the gut microbiome and host evolutionary history than between the gut microbiome and diet when looking at the level of host species. This phylogenetic signal of gut microbiome structure is well-supported in non-flying mammals and insects, but has been less well-supported in birds. Avian gut microbiome studies generally support differences between host species (Hird et al. 2015, San Juan et al. 2020, Capunitan et al. 2020, but see Hird et al. 2014), but phylosymbiosis was not supported in New Guinean passerines (Bodawatta et al. 2021a) and the signal was weak among captive cranes and in two passerine studies (Trevelline et al. 2020, Kropáčková et al. 2019, Loo et al. 2019). In the current study, concordance between the wood warbler phylogeny and gut microbiome dendrogram was moderate and similar to that reported for cranes in captivity (Trevelline et al. 2020) and passerines in the Czech Republic (Kropáčková et al. 2019). Thus, our results support the view that phylosymbiosis is weaker in birds than in mammals (Song et al. 2020, Youngblut et al. 2019) and uniquely demonstrate that in wood-warblers, a stronger influence of diet (as measured by species-level diet diversity) does not account for this discrepancy. Our findings of high variability of gut microbiomes for individuals within the same species may explain the lack of a consensus about phylosymbiosis in the avian literature, and particularly among studies that analyzed fewer individuals per species.

A phylogenetically conserved gut microbiome may provide the opportunity for co-adaptation between hosts and their gut microbes, which could implicate microbiomes in complex host evolutionary processes, including speciation (Brucker & Bordenstein 2012). Long-term coevolution between hosts and microbiota could explain phylosymbiosis, but this pattern could also arise under ecological filtering. Mazel et al. (2018) used simulations to show that under ecological filtering, the strength of phylosymbiosis is determined by the strength of the phylogenetic signal in the host trait underlying microbe colonization. It has been hypothesized that convergence of bat and avian gut microbiomes is due to reduced gut length, an adaptation to powered flight, which may favor rapid turnover in gut microbiota thus accounting for the weakened phylogenetic signal in gut microbiomes compared to non-flying mammals (Song et al. 2020). Consistent with this, Bodawatta et al. (2021a) found a negative association between passerine body mass—a proxy for gut length—and both gut microbiome richness and divergence. This might lead to the prediction that phylogenetic signal of the gut microbiome should be strongest in large-bodied birds, and weakest in small-bodied birds. However, the current data do not support this, as is highlighted by the results presented here. The strength of phylosymbiosis reported here for wood-warblers—small-bodied species weighing ∼6-20 g—is similar to that reported for cranes (Trevelline et al. 2020), which are several hundred times heavier. Thus, additional study is necessary to elucidate the effect of gut retention time on gut microbiome structure, and of other phylogenetically conserved avian traits or habitat preferences, including diet, that may mediate the colonization and maintenance of gut microbiomes.

Further study is also necessary to understand the biological relevance of taxonomic differences and of phylogenetic signal in gut microbiome structure between hosts. Experimental studies have shown antibiotic treatment administered to nestlings results in faster growth rates (Coates et al. 1963, Potti et al. 2002, Kohl et al. 2018), and caeca of germ-free chickens exhibit altered gene expression and notably do not express immunoglobulins (Volf et al. 2017). Thus, it is clear gut microbiota impose constraints on host development and immune function but how species-differences in natural gut microbiota composition might impact host fitness is unknown. It is important to note that although we observed an effect of host taxonomy on gut microbiome structure, this does not necessarily imply functional differences in gut microbiota between hosts. However, due to microbiome differentiation between host species we may predict disruption of these communities for admixed individuals upon hybridization (Brucker & Bordenstein 2012). Wood-warblers are well known to hybridize and occasionally even form intergeneric hybrids (Toews et al. 2018, 2020), making this clade an excellent system that can be used to tease apart evolutionary from ecological influences on the gut microbiome as well as the potential role of the microbiome in hybrid dysfunction.

### Caution against sequencing batch effects

We prepared and sequenced our 16S libraries in two different batches and the resulting yields were quite different (Figure S5). This strategy was desirable because it allowed us to process samples as they became available and it increased our overall sample sizes. However, we found the technical artifacts this introduced were not trivial (Figure S8), and similar to batch effects in other studies (Gibbons et al. 2018, Lou & Therkildsen 2022) it obscured the signal of the biological factors we tested (Table 2). Our topological congruence analysis seemed to be robust to the batch effect as our results were similar across datasets, although it is possible that batch effects obscured the signal of phylosymbiosis in previous avian gut microbiome studies. It is possible that batch effects are less of a concern for mammalian and other systems, where the signal of host phylogeny on gut microbiome structure is stronger than in birds. Similarly, the batch effect was less strong in the COI data which we also processed and sequenced in two batches (37% of total arthropod OTUs detected in both batches compared to 6% for bacterial ASVs). This may be due to a more rapid saturation of the accumulation curve for arthropod taxa than for bacterial taxa that are present in the warbler gut (Figure S2).

Future methodological study of the consequences of batch effects in metabarcoding studies is warranted. We recommend metabarcoding studies to report on sequences of technical replicates (PCRs amplified from the same sample within a batch) and positives (same sample sequenced across batches) which may help clarify when it is appropriate to make direct comparison of data sequenced in different batches.

### Concluding remarks

Our data highlight many outstanding questions about avian microbiomes and the ongoing need to characterize microbiomes of wild birds (Hird 2017). Wood-warbler gut microbiomes are dominated by Proteobacteria and Firmicutes, and on average, closely related host species share more similar gut microbiomes. We found little influence of sampling year, geographic locality, or diet diversity on gut microbiome structure and thus the majority of the variation between microbiota was left unexplained. Our results may suggest the phylogenetic signal in gut microbiome structure is tied more closely to host traits than to host ecology, yet the mechanisms driving this signal and possible functional consequences for hosts are not clear.

The level of phylogenetic signal in gut microbiome structure we detected is similar to that detected for larger-bodied birds (Trevelline et al. 2020), suggesting small body size does not preclude phylosymbiosis. Further study is necessary to understand the relationship between host body size, gut retention time, and gut microbe colonization. Although we found that broad-scale measures of diet diversity are not closely related to gut microbiome diversity, future studies should explore how components of the diet (e.g., dominant arthropod taxa, energetic values of food items) might influence the gut microbiome, including by way of their influence on host traits (e.g., gut pH). Wood-warblers represent a promising system to continue addressing outstanding ecological and evolutionary questions about the avian microbiome, including how microbiomes may influence and respond to adaptive radiation (Bodawatta et al. 2021b).

## Acknowledgements

We thank Brian Trevelline and Emily Davenport for helpful discussion and advice on data analyses, and we thank Laura Porturas and Lan-Nhi Phung for assistance in the field. This material is based upon work supported by the NSF Postdoctoral Research Fellowships in Biology Program under Grant No. 2010679. This work was also supported by the Department of Biology and the Huck Institute for Life Sciences at Penn State University, and by the Cornell Lab of Ornithology.

## Data accessibility and benefits-sharing

### Data accessibility

The sequencing data and metadata analyzed in this study will be deposited into the NCBI Short Read Archive upon acceptance of this manuscript. Scripts will be deposited to GitHub (github.com/baizm).

### Benefits-sharing

Benefits from this research accrue from the sharing of our data on public resources as described above.

## Author contributions

MDB and DPLT designed the study with input from all authors. DPLT, ETM, MDB, and AWW collected samples, and AWW processed samples and prepared sequencing libraries. All authors took part in data analysis. MDB and ABC wrote the manuscript, and all authors revised and approved the final version.

