## Supplementary material for "Gut microbiome composition better reflects host phylogeny than diet in breeding wood-warblers": Figure S1

### Host phylogeny, but not diet diversity, influences gut microbiome composition in breeding wood-warblers

#### Table of Contents:


| | |
|---|---|
| **Figure S1-S2** | Page 2 |
| **Figure S3-S4** | Page 3 |
| **Figure S5** | Page 4 |
| **Figure S6** | Page 5 |
| **Figure S7-S8** | Page 6 |
| **Figure S9** | Page 7 |
| **Figure S10** | Page 8 |
| **Figure S11** | Page 9 |
| **Table S1** | Page 10 |
| **Table S2** | Page 11 |
| **Table S3-S4** | File TableS3S4.xlsx |

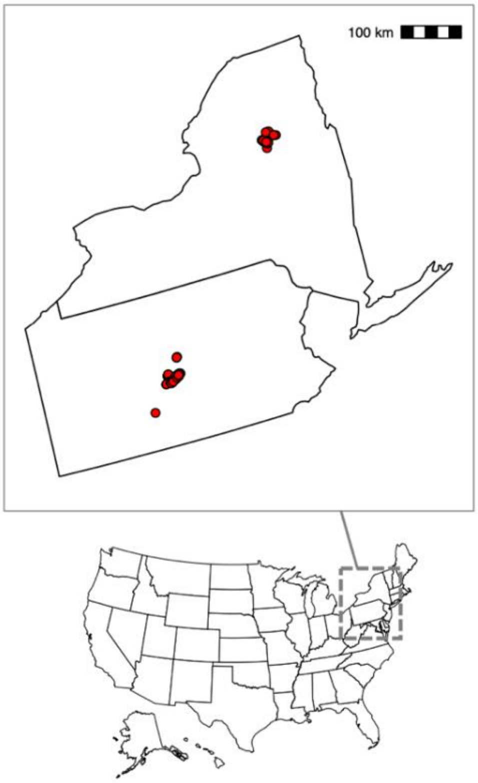


**Figure S1.** Map of sampling locations used in this study (red circles).

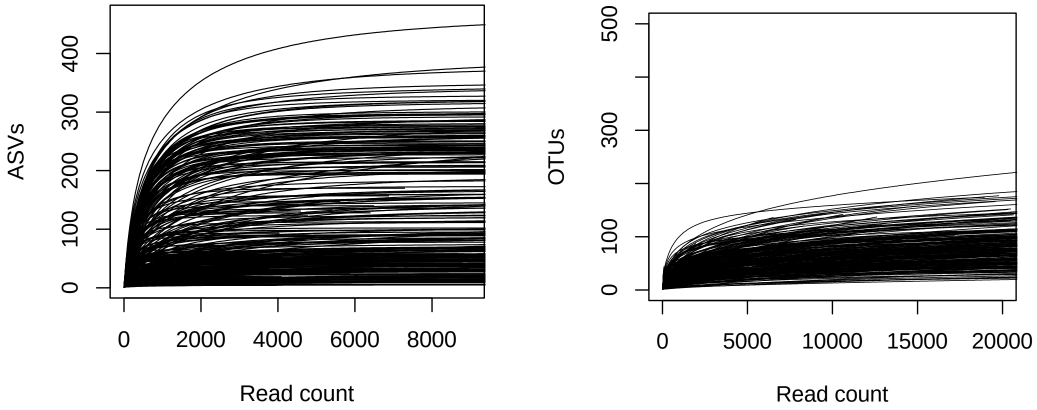


**Figure S2. A)** Rarefaction curve for 16S sequences, **B)** Rarefaction curve for COI sequences.

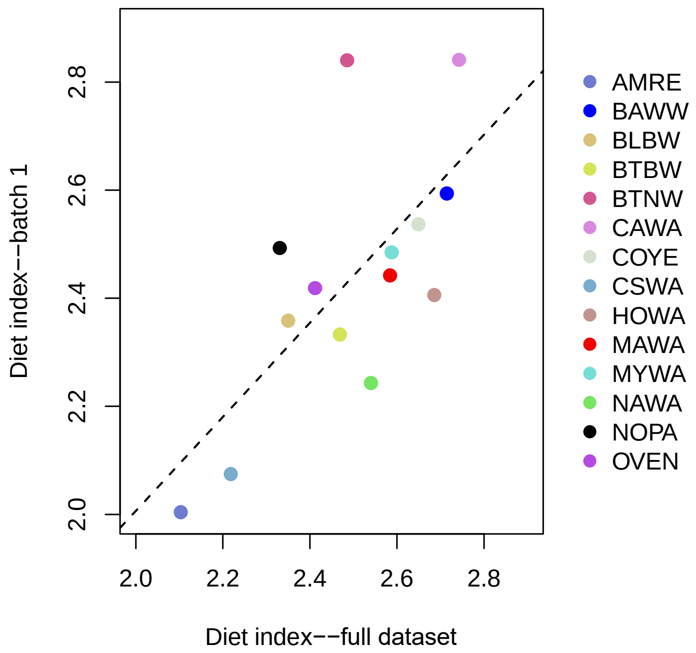


**Figure S3.** Species' diet index values are correlated between datasets (τ=0.516, *P*=0.010).

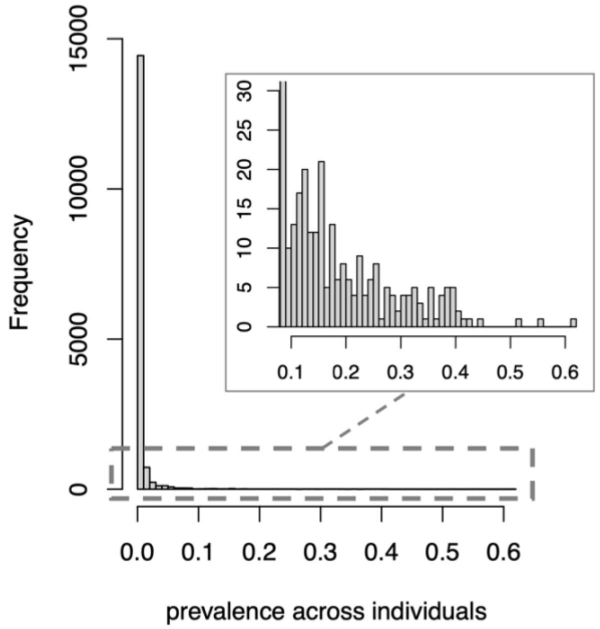


**Figure S4.** Frequency histogram of bacterial ASV prevalences across individuals in the full 16S dataset. ASV prevalence is calculated as proportion of individuals with ASV count > 0 (i.e., an ASV with a prevalence of 0.1 was present in 10% of the individuals in the study).

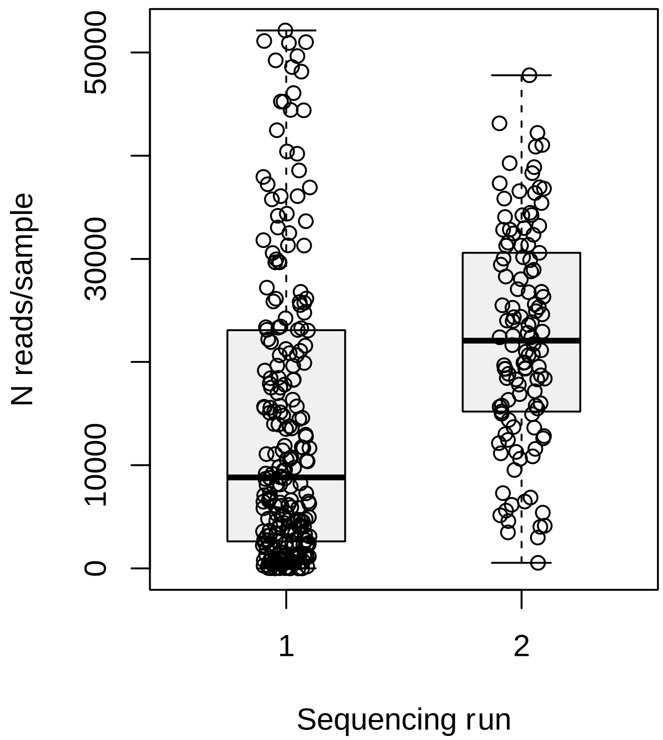


**Figure S5.** Raw number (N) of sequencing reads retrieved per sample across sequencing runs employed in this study. Sequencing run 1 contained samples collected between 2017-2019 (batch 1), and sequencing run 2 contained samples collected in 2020 (batch 2).

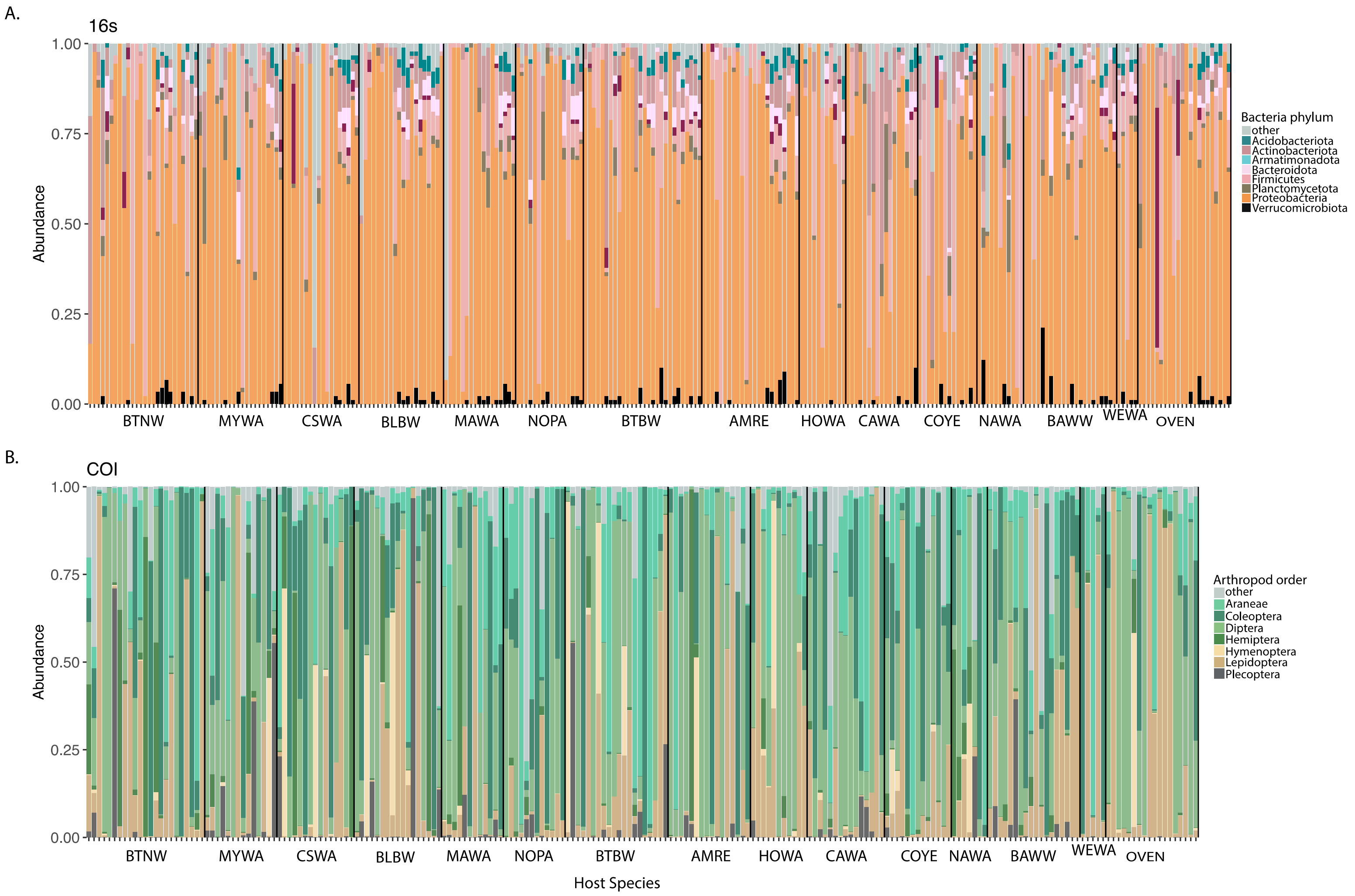


**Figure S6.** Individual-level relative abundance plots of **A)** top 8 bacterial phyla in the 16s full dataset, and **B)** top 7 arthropod orders in the COI full dataset.

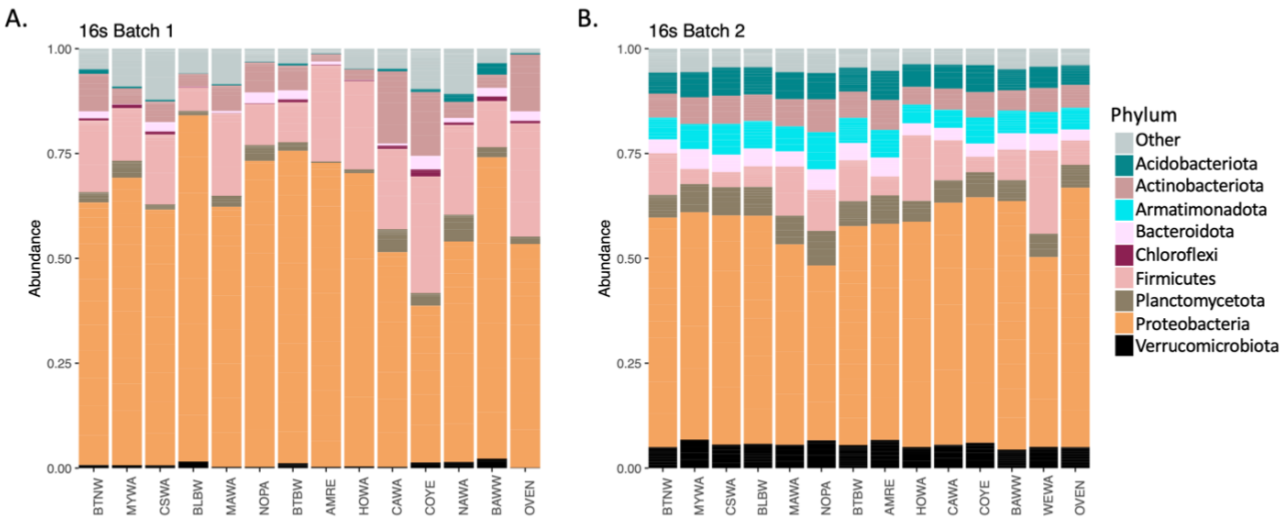


**Figure S7.** Relative abundance plots of top 8 bacterial phyla represented within each host species for **A)** samples collected between 2017-2019 that were sequenced in the first sequencing run, and **B)** samples collected in 2020 that were sequenced in the second sequencing run.

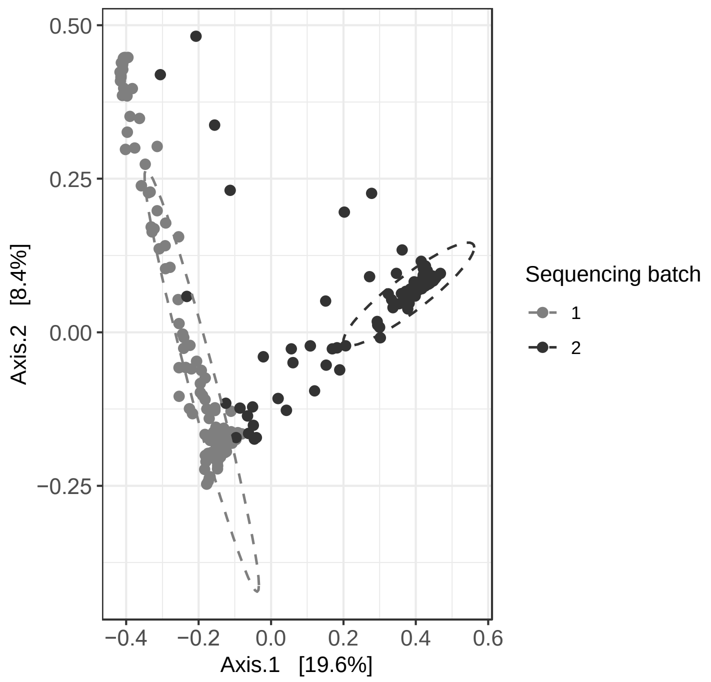


**Figure S8.** Principal coordinate analysis of Bray-Curtis distance between microbiomes in the full dataset, colored by sequencing run. Ellipses are drawn at 80% confidence level.

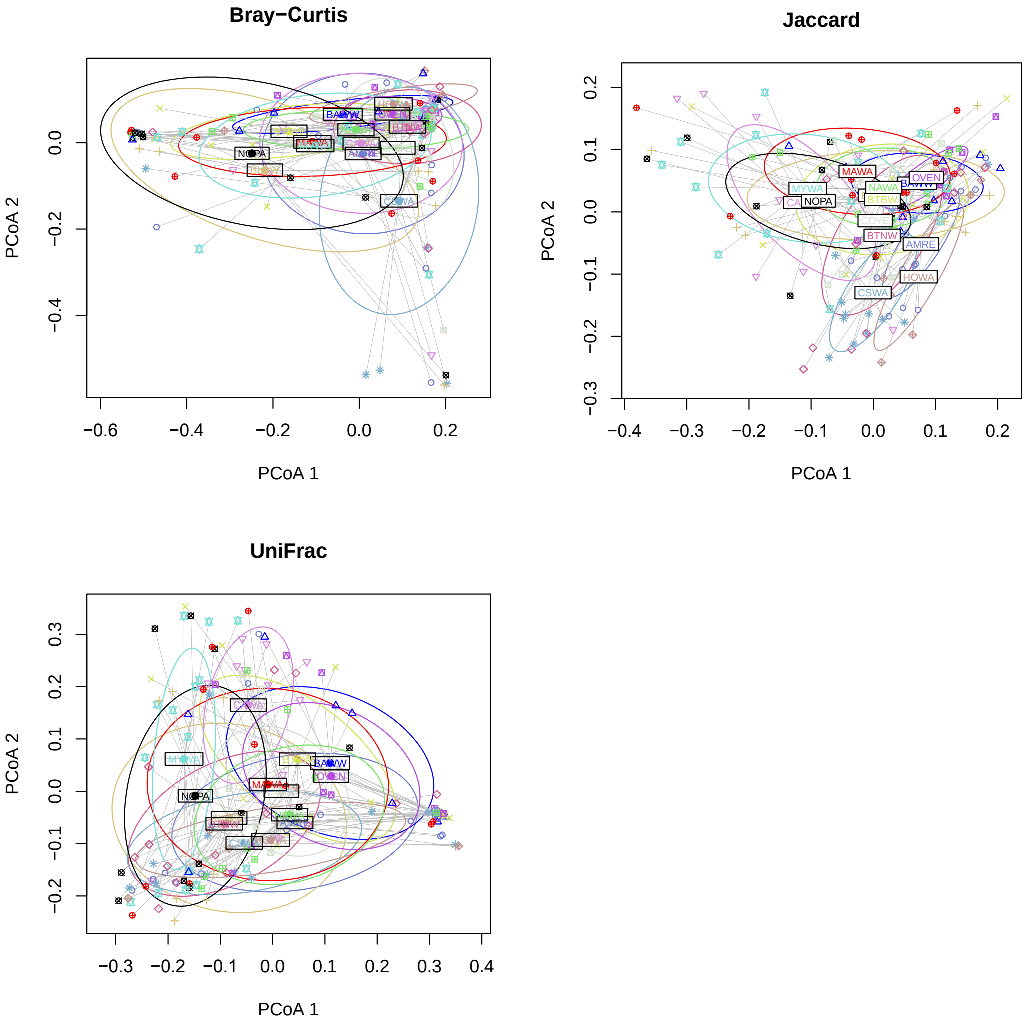


**Figure S9.** Multivariate dispersions of beta distances showing species' centroids (labeled in white box with species' codes) for **A)** Bray-Curtis distance, which exhibits homogenous dispersion (PERMUTEST F=0.703, *P*=0.762), **B)** Jaccard distance, which does not exhibit homogenous dispersion (PERMUTEST F= 2.409, *P*=0.009), and **C)** UniFrac distance, which does not exhibit homogenous dispersion (PERMUTEST F=3.049, *P*=0.002). Despite this heterogeneity in dispersion for Jaccard and UniFrac distances, species centroids are largely non-overlapping, as they are in the Bray-Curtis plot, suggestive of true differences between species in gut microbiome structure. Ellipses are 1 standard deviation from the centroid. See table 1 in the main text for host species codes.

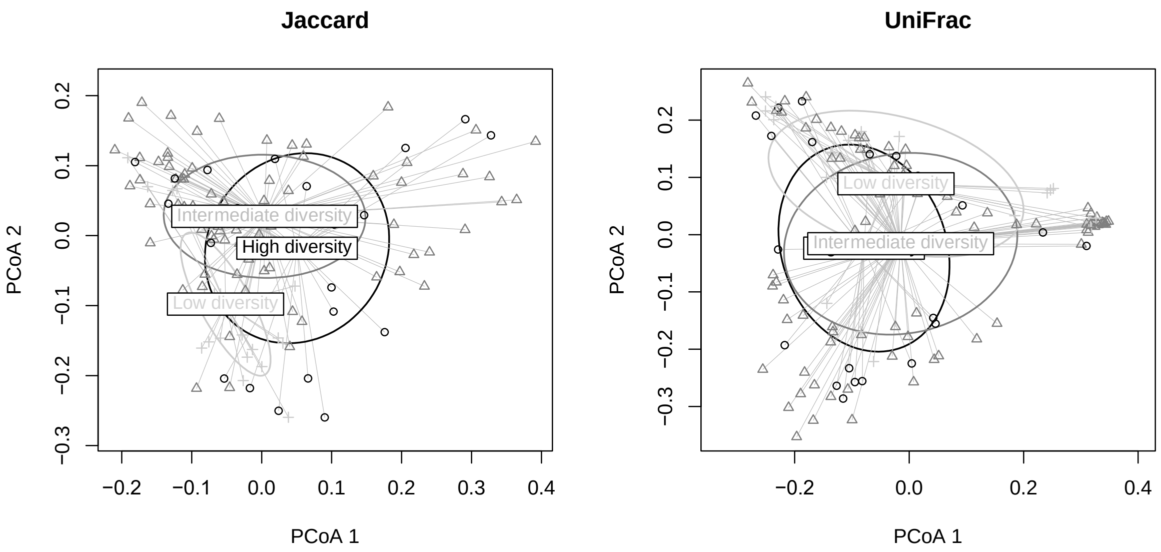


**Figure S10.** Multivariate dispersions of beta distances showing centroids among diet types (labeled in white box) for **A)** Jaccard distance, which does not exhibit homogenous dispersion (PERMUTEST F=9.067, *P*=0.001), and **B)** UniFrac distance, which does exhibit homogenous dispersion (PERMUTEST F=2.161 , *P*=0.118). In both plots diet type centroids are largely non-overlapping especially for low diversity diets. Ellipses are 1 standard deviation from the centroid.

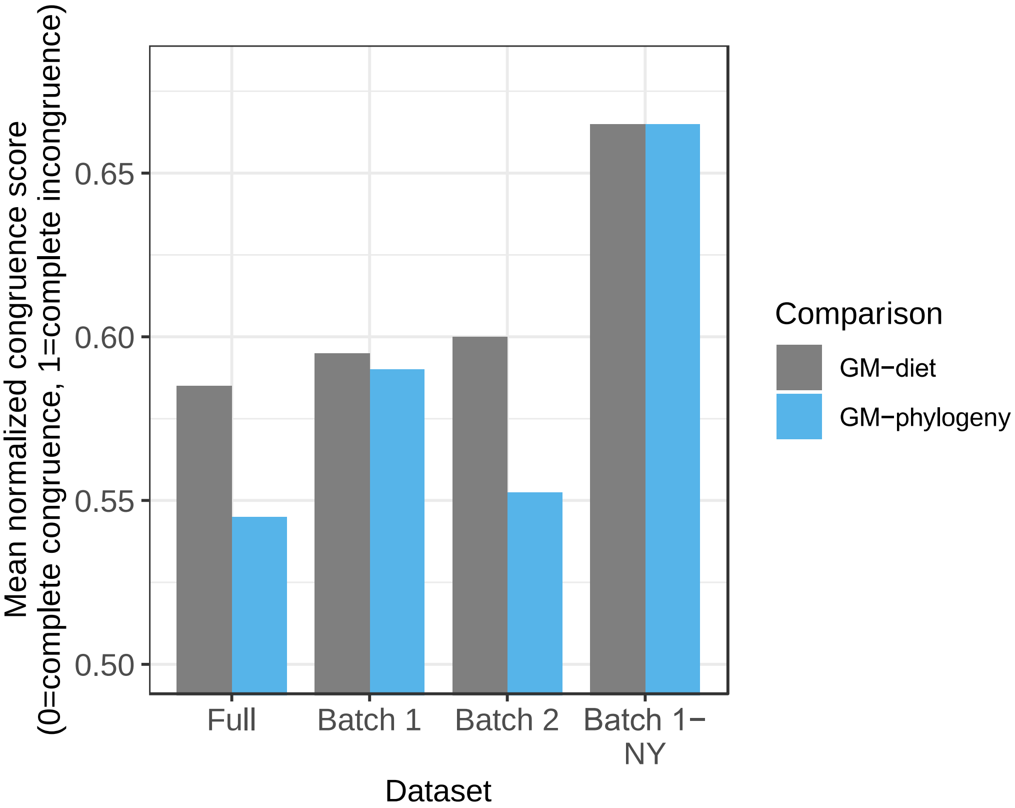


**Figure S11.** Mean congruence scores between the species-averaged gut microbiome (GM) dendrogram and the host species phylogeny (blue) and the species-averaged host diet dendrogram (gray). Each mean congruence score represents the mean across four analyses using Bray-Curtis, Jaccard, UniFrac, and Weighted UniFrac metrics, where low scores reflect high congruence. See Table 2 in the main text for statistical results of each test.

**Table S1.** Correlations between different diet diversity metrics. **A)** Alpha diversity of the diet. The Pearson correlation coefficient is reported. **B)** Beta diversity of the diet. The mantel statistic based on Spearman rank correlation is reported. In both panels, asterisks denote significance: ***p<0.001, **p<0.01, *p<0.05.

| **Table S1a. Diet alpha diversity** | | | |
|---|---|---|---|
| | | **Batch 1** | |
| | **Shannon index** | **Chao1** | **Faith's PD** |
| **Shannon index** | 1 | | |
| **Chao1** | 0.394*** | 1 | |
| **Faith's PD** | 0.704*** | 0.708*** | 1 |
| | | **Full dataset** | |
| | **Shannon index** | **Chao1** | **Faith's PD** |
| **Shannon index** | 1 | | |
| **Chao1** | 0.478*** | 1 | |
| **Faith's PD** | 0.520*** | 0.790*** | 1 |

| **Table S1b. Diet beta diversity** | | | | |
|---|---|---|---|---|
| | | **Batch 1** | | |
| | **Bray-Curtis** | **Jaccard** | **UniFrac** | **Weighted UniFrac** |
| **Bray-Curtis** | 1 | | | |
| **Jaccard** | 0.657** | 1 | | |
| **UniFrac** | 0.491** | 0.731** | 1 | |
| **Weighted UniFrac** | 0.377** | 0.199** | 0.129** | 1 |
| | | **Full dataset** | | |
| | **Bray-Curtis** | **Jaccard** | **UniFrac** | **Weighted UniFrac** |
| **Bray-Curtis** | 1 | | | |
| **Jaccard** | 0.622** | 1 | | |
| **UniFrac** | 0.308** | 0.635** | 1 | |
| **Weighted UniFrac** | 0.138** | 0.092* | 0.041 | 1 |

**Table S2.** Median divergence dates from TimeTree of Life used to scale the warbler phylogeny and derive cophenetic (evolutionary) distances between taxa. Nodes are from the topology in Figure 1a in the main text. MYA=million years ago.

| **Node** | **Divergence date (MYA)** |
|---|---|
| *Seiurus*-all taxa | 10.1 |
| *Cardellina-Setophaga* | 5.9 |
| *S. ruticilla* (AMRE)-*S. virens* (BTNW) | 5.3 |
| *Geothlypis-Leiothlypis* | 7.54 |
| *S. virens* (BTNW)-*S. coronata* (MYWA) | 5.3 |
| *S. pensylvanica* (CSWA)-*S. fusca* (BLBW) | 3.87 |
| *S. magnolia* (MAWA)-*S. americana* (NOPA) | 5.6 |
